# PDZD8-FKBP8 tethering complex at ER-mitochondria contact sites regulates mitochondrial complexity

**DOI:** 10.1101/2023.08.22.554218

**Authors:** Koki Nakamura, Saeko Aoyama-Ishiwatari, Takahiro Nagao, Mohammadreza Paaran, Christopher J. Obara, Yui Sakurai-Saito, Jake Johnston, Yudan Du, Shogo Suga, Masafumi Tsuboi, Makoto Nakakido, Kouhei Tsumoto, Yusuke Kishi, Yukiko Gotoh, Chulhwan Kwak, Hyun-Woo Rhee, Jeong Kon Seo, Hidetaka Kosako, Clint Potter, Bridget Carragher, Jennifer Lippincott-Schwartz, Franck Polleux, Yusuke Hirabayashi

**Author notes:** Corresponding author: Yusuke Hirabayashi, Ph.D., The University of Tokyo, Department of Chemistry and Biotechnology, School of Engineering 7-3-1 Hongo, Bunkyo-ku, Tokyo, Japan, 113-8656. Laboratory of Molecular Neurobiology, Institute for Quantitative Biosciences, The University of Tokyo, Tokyo 113-0032, Japan. Department of Neurosurgery, Stanford University School of Medicine, Stanford, CA 94304, USA. These authors contributed equally (Koki Nakamura, Saeko Aoyama-Ishiwatari, Takahiro Nagao).

## Abstract

Mitochondria-ER membrane contact sites (MERCS) represent a fundamental ultrastructural feature underlying unique biochemistry and physiology in eukaryotic cells. The ER protein PDZD8 is required for the formation of MERCS in many cell types, however, its tethering partner on the outer mitochondrial membrane (OMM) is currently unknown. Here we identified the OMM protein FKBP8 as the tethering partner of PDZD8 using a combination of unbiased proximity proteomics, CRISPR-Cas9 endogenous protein tagging, Cryo-Electron Microscopy (Cryo-EM) tomography, and correlative light-EM (CLEM). Single molecule tracking revealed highly dynamic diffusion properties of PDZD8 along the ER membrane with significant pauses and capture at MERCS. Overexpression of FKBP8 was sufficient to narrow the ER-OMM distance, whereas independent versus combined deletions of these two proteins demonstrated their interdependence for MERCS formation. Furthermore, PDZD8 enhances mitochondrial complexity in a FKBP8-dependent manner. Our results identify a novel ER-mitochondria tethering complex that regulates mitochondrial morphology in mammalian cells.

## Main

Mitochondria and the endoplasmic reticulum (ER) form contact sites (Mitochondria-ER contact sites: MERCS), an ultrastructural feature conserved in unicellular eukaryotes and metazoans. MERCS are the most abundant membrane contact sites (MCS) between organelles in many cell types and serve as a unique subcellular signaling platform for exchanging metabolites such as Ca^2+^ and glycerophospholipids. In addition to these critical biochemical reactions, key physiological and cell biological events essential for the maintenance of cellular homeostasis, including mitochondrial fission, mitochondrial DNA replication, and autophagosome biogenesis occur at these contact sites^1–3^.

Observations using electron microscopy (EM) have demonstrated that mitochondria and ER membranes are closely apposed at MCS, requiring proteins able to tether these two membranes within tens of nanometers of one another^4^. Intensive screening studies have identified multiple proteins localizing at MERCS in mammalian cells^1, 5–7^. Among those, the ER-resident protein PDZD8 was identified as a paralog of yeast Mmm1, a component of the ER–mitochondria encounter structure (ERMES)^8, 9^. Although the ERMES as a full complex formed by four proteins is lost in mammals, PDZD8 is required for forming the majority (∼40-80%) of MERCS in various cell types, and its deletion in cell lines and in mammalian neurons results in the disruption of intracellular Ca^2+^ dynamics by decreasing the fraction of Ca^2+^ released from the ER that can be imported directly into mitochondria^8, 10–12^. Consistent with its role in neurons of central nervous system (CNS), PDZD8 regulates dendritic Ca^2+^ dynamics in hippocampal CA1 and underlies their response properties *in vivo*^13^. Furthermore, genetic loss of function mutations of *PDZD8* in humans leads to syndromic intellectual disability^14^. In addition, expression quantitative trait loci (eQTL) mapping identified a single-nucleotide polymorphism affecting the expression of PDZD8 in the dorsolateral prefrontal cortex in a population of patients with high risk for post-traumatic stress disorder (PTSD)^15^. Therefore, PDZD8 plays a critical role in controlling neuronal and circuit function, and proper brain development and homeostasis in mammals^12^.

In addition to MERCS, the ER forms various MCSs with other organelles such as lysosomes, endosomes, Golgi apparatus, lipid droplets, and the plasma membrane (PM)^2, 16–19^. We and other groups have previously shown that PDZD8 localizes at MERCS in various cell types^8, 20, 21^. However, it has also been reported recently that overexpression of Rab7 or LAMP1 can recruit PDZD8 to the ER-late endosome or ER-lysosome contact sites, respectively^20, 22, 23^. In addition, overexpression of PDZD8 and Rab7 recruits the mitochondria to ER-endosome contact sites and was proposed to lead to the formation of three-way MCS^20^. Therefore, PDZD8 might participate in the formation of MCS networks besides tethering MERCS. As such, we hypothesized the existence of a currently unknown molecular effector required to recruit PDZD8 specifically to MERCS. To elucidate the molecular mechanisms underlying PDZD8-dependent MERCS formation, we used multiple independent proximity-based proteomic approaches relying on endogenous protein tagging. Since overexpression of PDZD8 can alter its subcellular distribution^8^. we implemented CRISPR-Cas9 technology to generate knock-in cell lines where endogenous PDZD8 is tagged with various epitopes, fluorescent proteins or catalytic enzymes, allowing its localization by microscopy or proximity-based proteomic screens. We demonstrate that the mitochondrial LC3 receptor FK506 binding protein 8 (FKBP8 also known as FKBP38) is a novel, direct PDZD8-interacting protein, and that the PDZD8-FKBP8 complex is required for MERCS formation in metazoan cells. Using combinations of Cryo-EM tomography and correlative light-electron microscopy (CLEM), we revealed the ultrastructural features of MERCS mediated by the PDZD8-FKBP8 tethering complex. Finally, our serial scanning electron microscopy demonstrated that PDZD8 regulates mitochondrial complexity through inhibition of FKBP8 function.

## Results

### Subcellular localization of endogenous PDZD8

While we previously reported that a significant fraction of PDZD8 localizes at MERCS^8^, PDZD8 was recently shown to localize to the ER-late endosome and ER-lysosome contact sites^11, 20, 22, 23^. Some of these studies failed to detect the enrichment of PDZD8 at MERCS, however, in the absence of reliable antibodies detecting endogenous PDZD8 protein by immunofluorescence, these studies often relied on overexpression of tagged forms of PDZD8 which disrupts both its subcellular localization and can generate gain-of-function phenotypes, for instance by increasing the number and size of MERCS or other MCSs where it is localized. Therefore, to determine the subcellular distribution of endogenous PDZD8 protein at MCS formed by the ER, we developed a knock-in mouse embryonic fibroblast NIH3T3 cell line fusing the fluorescent protein Venus sequence to the C-terminus of the *Pdzd8* coding sequence (**Extended Data Fig. 1a, b**). To avoid an artifactual increase in size and/or biogenesis of late endosome/lysosome due to overexpression of key effector proteins Rab7 or LAMP1, colocalization analyses were performed by detecting these proteins at endogenous levels with antibodies against endogenous markers: LAMP1 for lysosomes, Rab7 for the late endosomes, and Tomm20 and OXPHOS proteins for mitochondria. In agreement with previous studies, confocal microscopy imaging showed that 14.8% of PDZD8-Venus visualized by an anti-GFP antibody overlapped with LAMP1 staining, and under these endogenous expression conditions 7.7% overlapped with Rab7 staining. However, a significantly larger fraction overlapped mitochondria labeled either with Tomm20 (25.0%) or OXPHOS staining (22.1%), suggesting that endogenous PDZD8 is present at multiple MCS but is most abundant at MERCS (**Extended Data Fig. 1c,d**).

We next investigated the dynamics of endogenously expressed PDZD8 using time-lapse imaging in live cells. The *PDZD8*-HaloTag KI HeLa cell line (**Extended Data Fig. 2a, b**) was transiently transfected with the ER-localized reporter (BiP-mtagBFP2-KDEL) and an outer mitochondrial membrane (OMM)-localized reporter (YFP-ActA^24, 25^). The PDZD8-HaloTag was labeled with Janelia Fluor 549 dye. Triple-color time-lapse imaging using confocal microscopy demonstrated that PDZD8-Halotag puncta can be stably localized at MERCS despite significant dynamics of both ER and mitochondria, suggesting a direct association of PDZD8 with mitochondria may be present (**Extended Data Fig. 2c, Supplementary video 1**).

### Single molecule imaging of PDZD8 at the membrane contact sites

Recent work demonstrated that the ER-resident MERCS forming protein VAPB exhibits transient but highly frequent visits to MERCS. Thus, to determine the localization and molecular dynamics of PDZD8 along the ER membrane relative to MERCS or other MCS, we performed single particle tracking-photoactivation localization microscopy (sptPALM)^26^. Single PDZD8 molecules were visualized by labeling overexpressed PDZD8-HaloTag with a photoactivatable version of JF646 in COS7 cells (**Supplementary video 2**). Analysis of localization probabilities using a spatially defined probability function^27^ revealed PDZD8 localization was entirely restricted to the ER **(Fig. 1a)**. Strikingly, we observed regions along the ER where the probability was significantly higher (hotspots), presumably as a result of tethering and engagement with interacting proteins at contact sites with other organelles. In agreement with our endogenous labeling, ∼47% of these hotspots were in close proximity with mitochondria **(Fig. 1a–c**, Extended Data Fig. 3; 90 out of 192 hotspots). By following the trajectories of single PDZD8 molecules outside and within these putative mitochondria-contact sites (MitoCS), we found that PDZD8 can dynamically enter and exit these hotspots in seconds (**Fig. 1B and C, Supplementary video 3**). Importantly, the effective diffusion (D_eff_) of single PDZD8 molecules within MitoCS was significantly reduced compared the rest of the ER (0.22 ± 0.0025 μm^2^/s in MitoCS, mean ± SEM, n = 90; **Fig. 1e**) suggesting that PDZD8 is captured at MERCS but still remains mobile at these contact sites. Consistent with this, PDZD8 single particles dwell at the hotspots for a median time of just 1.1 seconds per each visit **(Fig. 1g)**. In addition to MitoCS, we also observed spots with high probability of PDZD8 that were not mitochondria-associated, which might correspond to non-mitochondrial organelle-contact sites (OtherCS, **Fig. 1a,b,d**). The D_eff_ and dwell time of PDZD8 in the OtherCS is similar to those in MitoCS, but the mean of individual CS area was significantly larger at MitoCS than in the OtherCS **(Fig. 1f).**

**Figure 1.**
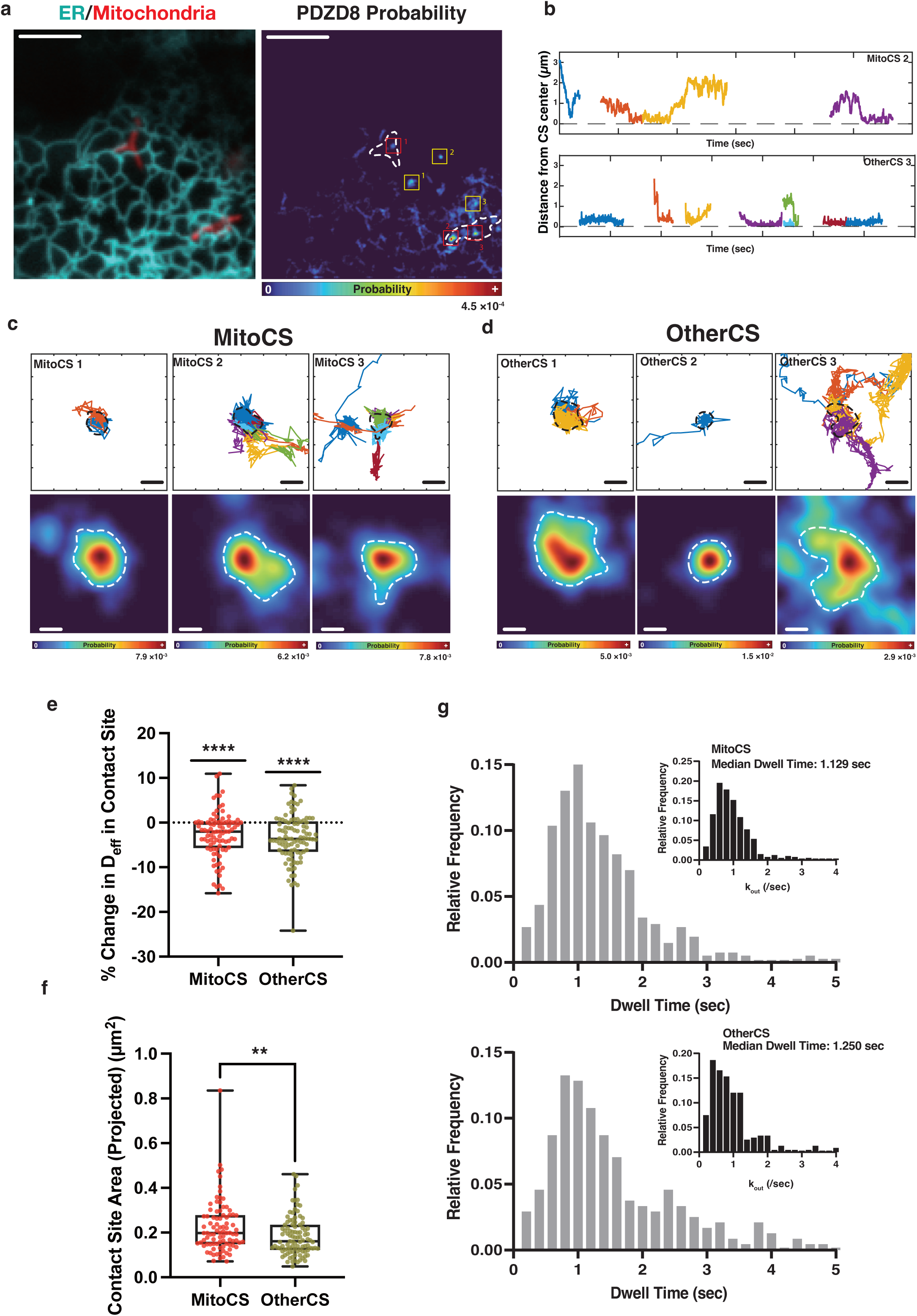
PDZD8 shows specific interactions at both mitochondria and at other organelles in the same cells by high-speed single molecule tracking. **a,** Diffraction-limited imaging of the ER (cyan) and the mitochondria (red) in the periphery of a representative COS7 cell with the simultaneously measured likelihood of finding a PDZD8 molecule in a one-minute window. The locations of the mitochondria in the probability map are indicated with dotted white lines. Boxes correspond to the mitochondria-(red) or non-mitochondria-(yellow) contact sites in **c** and **d**. **b**, Plots of the distance of individual PDZD8 molecules shown in different colors from the center of example contact sites over time. Plots are from MitoCS 2 (top panel) or OtherCS CS3 (bottom panel) shown in **a**. Note the long stretches where single molecules remain engaged in the contact site. **c,** Zooms of the MitoCS indicated in **a** showing individual PDZD8 trajectories engaging with the contact site and the associated PDZD8 probability density. Dotted lines indicate contact site boundaries as used for subsequent analysis. **d,** Zooms of OtherCS in **a**, dotted lines represent the contact site boundary as used in subsequent analysis. **e,** PDZD8 shows reduced diffusion within both classes of contact sites as compared to freely diffusing in the surrounding ER. **f,** Sizes of MitoCS are significantly larger than those of OtherCS in the same cells. **g,** PDZD8 dwell times in individual MitoCS and OtherCS. Inset shows the leaving frequency (k_out_) of individual PDZD8 molecules from the MitoCS and OtherCS.

The molecular dynamics of PDZD8 at MERCS are different than the ER proteins VAPA or STIM1 localizing at ER contact sites with the plasma membrane^28, 29^. We note that despite the rapid tether exchange reported here, the size of the contact sites observed with PDZD8 is significantly larger and the mean dwell time of PDZD8 at the contact site was significantly longer compared to those reported with VAPB **(Fig. 1f, g)**^30^. The fact that the behavior of PDZD8 and VAPB at the MCS differs suggests that PDZD8 interaction at MERCS may represent a distinct, specific tethering from the interactions described for VAPB. Taken together, these data suggest that PDZD8 is highly dynamic along the ER but drastically slows down at contacts between ER and mitochondria as well as other potential MCS between ER and other organelles. These results strongly suggest the existence of an unknown tethering partner for PDZD8 along the OMM.

### Identification of FKBP8 as a binding partner of PDZD8 using unbiased *in vivo* proteomic screens

To identify the tethering partner of PDZD8 facilitating this behavior at MERCS, we designed unbiased proteomic screens using endogenous PDZD8 protein immunoprecipitation coupled with mass spectrometry (IP-MS) (**Fig. 2a**). To avoid artifacts due to PDZD8 overexpression, we established a new mouse line engineered with a 3×HA tag fused to the endogenous PDZD8 protein using CRISPR-Cas9 mediated genomic knock-in (*Pdzd8*-3×HA KI mouse line) (**Fig. 2b**, Extended Data Fig. 4a, b). Since PDZD8 is expressed at high levels in neurons, protein complexes containing PDZD8 were isolated from the neocortex of either *Pdzd8*-3×HA KI mice or control littermates at postnatal 10 days by IP using anti-HA antibody. Identification of the corresponding proteins immunoprecipitated in complex with PDZD8-3×HA by LC-MS/MS revealed that, in addition to previously identified PDZD8 interactors such as Protrudin, VAPA and VAPB^11, 31^, proteins known to localize at MERCS and/or mitochondria were significantly enriched in the immunoprecipitates from the KI mice compared to the control mice (**Fig. 2c, Table S1**).

**Figure 2.**
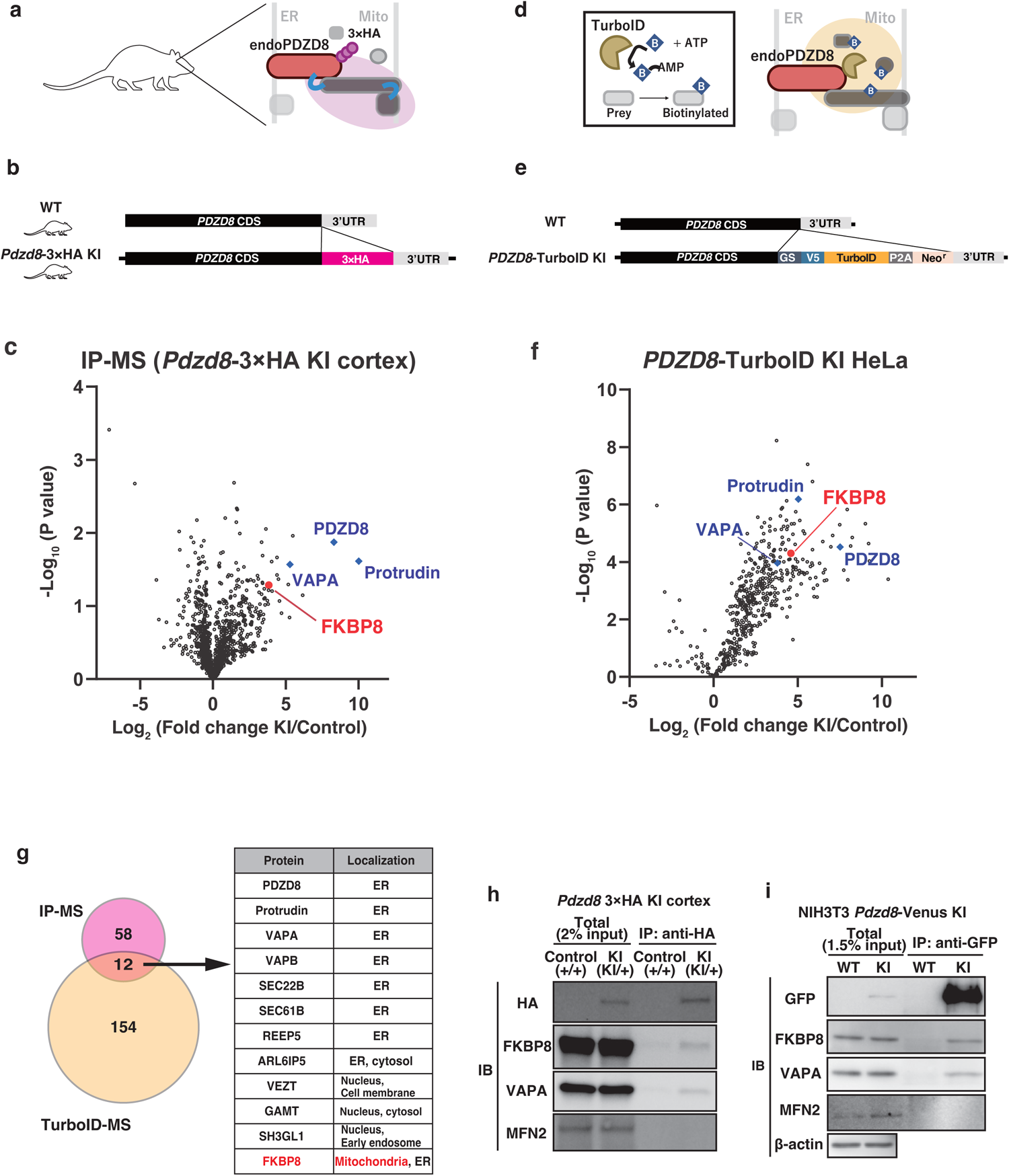
Proteomics screening identified a PDZD8-FKBP8 protein complex. **a,** Scheme of the immunoprecipitation and LC-MS/MS analysis using the *Pdzd8*-3×HA KI mice neocortices. The immunoprecipitates from the neocortices of *Pdzd8*-3×HA mice or the control littermates using an anti-HA antibody were subjected to the LC-MS/MS analysis. **b,** Diagram describing the genomic sequence of *Pdzd8*-3×HA KI mice. The sequence of a 3×HA tag was knocked-in at the C-terminus of the *Pdzd8* coding sequence. **c,** Volcano plot of proteins differentially binding to the PDZD8-3×HA. FKBP8 is labeled in red. Protrudin and VAPA, which have been previously reported to interact with PDZD8, are labeled in blue. The plot represents data from three biological replicates. **d,** Scheme of labeling proteins in the vicinity of endogenous PDZD8. A Biotin ligase TurboID fused to PDZD8 generates biotin–5ʹ-AMP from biotin and ATP. The biotin–5ʹ-AMP can covalently bind to proteins located within about 20 nm of endogenously expressed PDZD8-TurboID. After tryptic digestion, the biotinylated peptides were enriched using tamavidin 2-REV beads and subjected to the LC-MS/MS analysis. **e**, Diagram describing the genomic sequence of the *PDZD8*-TurboID KI HeLa cell. The sequence of TurboID-P2A-Neo^r^ was knocked-in at the C-terminus of the *PDZD8* coding sequence. **f,** Volcano plot of proteins differentially biotinylated with biotin in the *PDZD8*-TurboID KI HeLa cell. FKBP8 is labeled in red. Protrudin and VAPA, which have been previously reported to interact with PDZD8, are labeled in blue. The volcano plot represents three biological replicates. **g,** Numbers of proteins highly enriched in the IP-MS (**c**) and TurboID-MS (**f**) are shown in a Venn diagram. The highly enriched proteins were selected according to the following criteria: log_2_ (fold change) > 1 and -log_10_ (p-value) > 1 for IP-MS and log_2_ (fold change) > 3, -log_10_ (p-value) > 1.25 for TurboID-MS. 12 proteins are commonly found in the two proteomes. Note that FKBP8 is the only protein annotated with mitochondrial localization. **h-i**, Analysis of the interaction between endogenous FKBP8 and endogenous PDZD8-3×HA from the mouse neocortex (**h**), or endogenous PDZD8-Venus from NIH3T3 cells. Extracts from neocortex in *Pdzd8*-3×HA KI mouse (**h**) or *Pdzd8*-Venus KI NIH3T3 cells (**i**) were subjected to immunoprecipitation (IP) with antibodies to HA or GFP respectively. The resulting precipitates as well as the original tissue extracts (Total) were subjected to immunoblot analysis with antibodies to FKBP8, VAPA, MFN2, HA (**h**), GFP (**i**), and β-actin (**i**).

Next, in order to narrow down the protein list to only proteins in close proximity to PDZD8, we employed an independent approach, a proximity-based labeling screen using a biotin ligase TurboID (**Fig. 2d**)^32^. Again, to avoid overexpression-induced artifacts, we established a *PDZD8*-TurboID KI HeLa cell line using CRISPR-Cas9 knock-in technology (**Fig. 2e**). These *PDZD8*-TurboID KI HeLa cells were treated with biotin for 6 hours and biotinylated peptides were isolated using tamavidin 2-REV beads and identified by LC-MS/MS (**Fig. 2f, Table S2**). Among 166 proteins identified by this screening approach, 12 proteins were also identified by the IP-MS-based screen (**Fig. 2g**). Among these candidate interactors, the only protein previously shown to localize at the outer mitochondrial membrane (OMM) was FKBP8 (**Fig. 2g**). Finally, we also performed a proteomic screen using TurboID in a mouse neuroblastoma cell line (Neuro2a) and again identified FKBP8 in the list of biotinylated proteins (Extended Data Fig. 4c, d). Specific co-immunoprecipitation of FKBP8 and PDZD8 was confirmed by Western blotting using the *Pdzd8*-3×HA knock-in mouse (**Fig. 2h**) and *Pdzd8*-Venus KI NIH3T3 cells (**Fig. 2i**). These three independent proteomic approaches converge to strongly suggest that PDZD8 and FKBP8 reside in the same protein complex.

### Direct interaction between PDZD8 and FKBP8 proteins

To test if the interaction between PDZD8 and FKBP8 is direct, we measured the binding affinity of PDZD8-FKBP8 interaction *in vitro.* We used surface plasmon resonance (SPR) with purified recombinant cytosolic portions of both FKBP8 and PDZD8 (**Extended Data Fig. 5a, b).** Recombinant PDZD8 proteins without its transmembrane domain (ΔTM) were immobilized on a sensor chip and changes of the surface resonance upon recombinant FKBP8ΔTM injection were measured. As expected, the SPR responded in a FKBP8 dose-dependent manner in the 2 to 90 µM range (**Fig. 3a**). Fitting R_eq_ (SPR responses in equilibrium) and FKBP8 concentration to the monovalent binding model provided K_D_ value of 142 µM (**Fig. 3b**). Thus, the affinity between recombinant PDZD8 and FKBP8 is in the same range as other previously reported VAPB-PTPIP51 MERCS tethering complex and agrees with the unexpectedly rapid exchange observed by sptPALM^33^.

**Figure 3.**
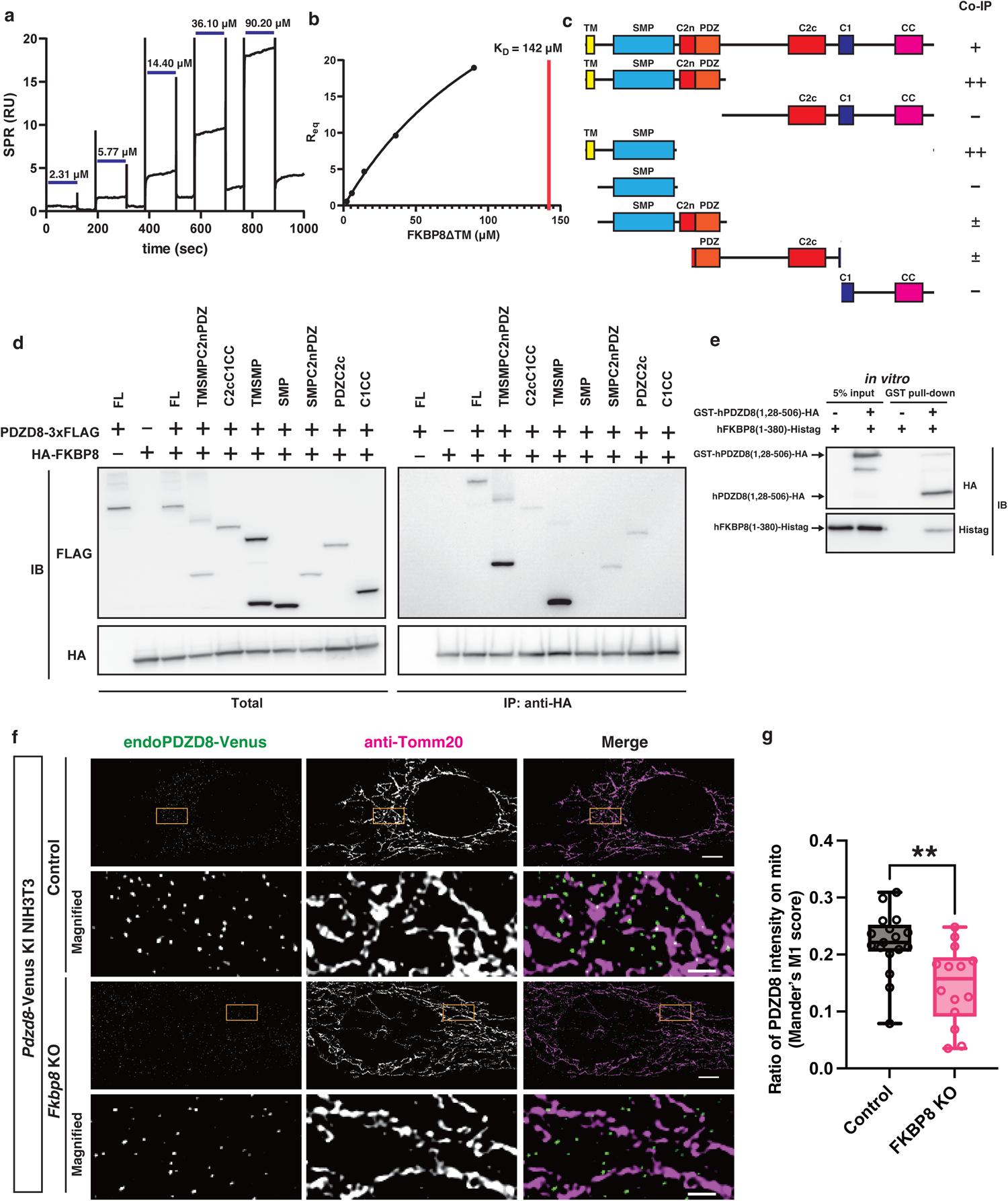
A direct binding partner FKBP8 is required for PDZD8 to be recruited on mitochondria. **a,** Sensorgrams of SPR assay. Recombinant human PDZD8 (1,28-) – FLAG was immobilized to the sensor chip and FKBP8 (1-380) – Histag with indicated concentrations were injected. **b,** SPR responses in the equilibrium (R_eq_) are plotted against FKBP8 concentration. The plot of R_eq_ versus FKBP8 concentration was fitted to the monovalent binding model to determine K_D_ values. **c,** Schematic diagram of the mutants of PDZD8 deleted with various domains. TM; Transmembrane, SMP; Synaptotagmin-like mitochondrial-lipid-binding, C2n; N-terminal sequence of C2 domain, PDZ; PDZ domain, C2c; C-terminal sequence of C2 domain, C1; C1 domain, CC; coiled-coil region. **d,** *Pdzd8*^f/f^::Cre^ERT2^ MEFs expressing a series of deletion mutants of PDZD8-3×FLAG shown in **c** and HA-FKBP8 were treated with 1 μM 4-hydroxy tamoxifen (4-OHT) and cell extracts were immunoprecipitated with anti-HA antibody. Western Blotting was performed with anti-HA antibody and anti-FLAG antibody. **e**, GST-Pulldown assay from the mixture of recombinant GST - Thrombin cleavage site - human PDZD8 (1, 28-50–) - HA and recombinant human FKBP8 (1-38–) - Histag *in vitro*. FKB–8 - Histag was isolated only with the GST-beads incubated with GST-PDZD8 (1, 28-506) – HA. **f,** Immunofluorescence analysis of *Pdzd8*-Venus KI NIH3T3 cells knocking out endogenous FKBP8 by confocal microscopy with a Nikon Spatial Array Confocal (NSPARC) detector. The cells were transfected with the control gRNA (upper two rows) or three gRNAs against FKBP8 (bottom two rows), Cas9, and transfection marker mtagBFP2, and stained with antibodies to GFP, and Tomm20 for visualizing endogenous PDZD8-Venus (green) and mitochondrial outer membrane (magenta), respectively. Scale bar: 5 µm (original) 1 µm (magnified). **g,** Quantification of the percentage of endogenous PDZD8-Venus intensity overlapping with mitochondria (Tomm20-positive area).n = 17, 14 cells for the control and FKBP8 KO cells. Statistical analysis was performed using Mann-Whittney U test. **p < 0.01

### TM and SMP domains of PDZD8 are sufficient for FKBP8 binding

Next, to identify the protein domains of PDZD8 required to mediate interaction with FKBP8, we conducted co-IP experiments by expressing a series of 3×FLAG-tagged PDZD8 deletion mutants together with HA-tagged FKBP8 (**Fig. 3c**). Based on the previous reports suggesting that PDZD8 can homodimerize, endogenous full-length PDZD8 can act as a bridge between exogenously expressed truncated forms of PDZD8 and FKBP8, even in the absence of direct binding^23^. To avoid this, we established a tamoxifen-inducible *Pdzd8* conditional KO mouse embryonic fibroblast cell line (*Pdzd8*^f/f^::Cre^ERT2^ MEFs) (**Extended Data Fig. 5c-e**). Using a time course analysis, we determined that PDZD8 was undetectable 45 hours after Cre^ERT2^-mediated deletion of the floxed allele by treatment with 4-hydroxytamoxifen (4-OHT; **Extended Data Fig. 5f**). Whereas truncated forms of PDZD8 including TM-SMP domains co-precipitated FKBP8 efficiently, none of the other domains showed strong binding to overexpressed FKBP8 (**Fig. 3d**). These results suggest that TM-SMP domains of PDZD8 represent the minimal domain mediating interaction with FKBP8. Given that previous protein binding and late endosome/lysosome recruitment functions of PDZD8 are independent of the SMP domain^20, 22, 34^, the SMP domain of PDZD8 represents a unique binding interface with FKBP8.

Next, we tested if SMP-C2n-PDZ domain of PDZD8 directly binds to FKBP8 using purified recombinant proteins. Recombinant glutathione S-transferase (GST) - Thrombin cleavage site - human PDZD8 (1,28- 506) - HA and human FKBP8 (1-380) - Histag were expressed in *E.coli.* and purified with GST-binding beads and TALON affinity columns, respectively. These purified proteins were mixed *in vitro*, applied to a column with GST-binding beads and eluted by cleaving the thrombin cleavage site. Western blotting analysis revealed that FKBP8 was isolated only when it was incubated with GST-PDZD8 (1,28-506) - HA (**Fig. 3e**). Collectively, these results demonstrate that FKBP8 is a direct binding partner of PDZD8 through the SMP-C2n-PDZ domain.

### FKBP8 is required for the recruitment of endogenous PDZD8 to mitochondria

Our live imaging of endogenously expressed PDZD8 and the single molecule tracking of PDZD8 showed that PDZD8 is highly mobile throughout the ER but shows distinct interactions (confined diffusion) where the ER is contacting mitochondria (Extended Data Fig. 3, **Fig. 1e, f, Supplementary video 1).** Therefore, the direct binding of PDZD8 and FKBP8 prompted us to examine whether FKBP8 is required for capturing of PDZD8 to mitochondria. To achieve this, Cas9 and guide RNAs targeting Fkbp8 gene locus were transiently expressed in the PDZD8-Venus KI NIH3T3 cell line. Immunocytochemistry confirmed that FKBP8 was not detectable in more than 81% of cells transfected (labeled by mTagBFP2; **Extended Data Fig. 6a**). We quantified the ratio of PDZD8 closely associated with mitochondria (stained by the anti-Tomm20 antibody) using a confocal microscopy equipped with a super resolution Nikon Spatial Array Confocal (NSPARC) detector. While mitochondrial areas per cells were not significantly affected by FKBP8 depletion, colocalization of PDZD8 on the mitochondria was significantly reduced in the FKBP8-depleted cells. **(Fig. 3f, g).** This demonstrate that FKBP8 is required for recruiting the ER protein PDZD8 to mitochondria.

### PDZD8 and FKBP8 are cooperatively required for formation of MERCS

Our results demonstrate that a direct binding between FKBP8 and PDZD8 and also that FKBP8 is required for PDZD8 recruitment to mitochondria. Thus, we investigated if the interaction between PDZD8 and FKBP8 is critical for MERCS formation. To measure the size of MERCS, we visualized the ER and mitochondria membranes using scanning electron microscopy (SEM) and segmented the contact sites between the two organelles. As previously reported in HeLa cells constitutively deleted with PDZD8^8^, conditional deletion of PDZD8 induced by a treatment with 4-OHT to *Pdzd8*^f/f^::Cre^ERT2^ MEFs (*Pdzd8* cKO) significantly decreased the size of MERCS, defined as the fraction of OMM membranes associated (<23.4nm) with ER, compared to the vehicle-treated control isogenic MEFs (**Fig. 4a, b**). Strikingly, shRNA mediated knock-down (KD) of *Fkbp8* (validated in **Extended Data Fig. 6b**) in the vehicle-treated control MEFs significantly decreased the size of MERCS to the same extent as in conditional *Pdzd8* cKO cells (**Fig. 4a, b**). Importantly, *Fkbp8* KD in *Pdzd8* cKO MEFs did not further reduce the fraction of MERCS compared to *Pdzd8* KO MEF only (**Fig. 4a, b**, *Pdzd8* cKO vs *Pdzd8* cKO + *Fkbp8* KD). Two-way ANOVA analysis shows that there is a strong functional interaction between the effects of FKBP8 and PDZD8 loss of function regarding the size of MERCS (**Fig. 4c, d**). Therefore, these results demonstrate that PDZD8 and FKBP8 tether the ER and mitochondria interdependently.

**Figure 4.**
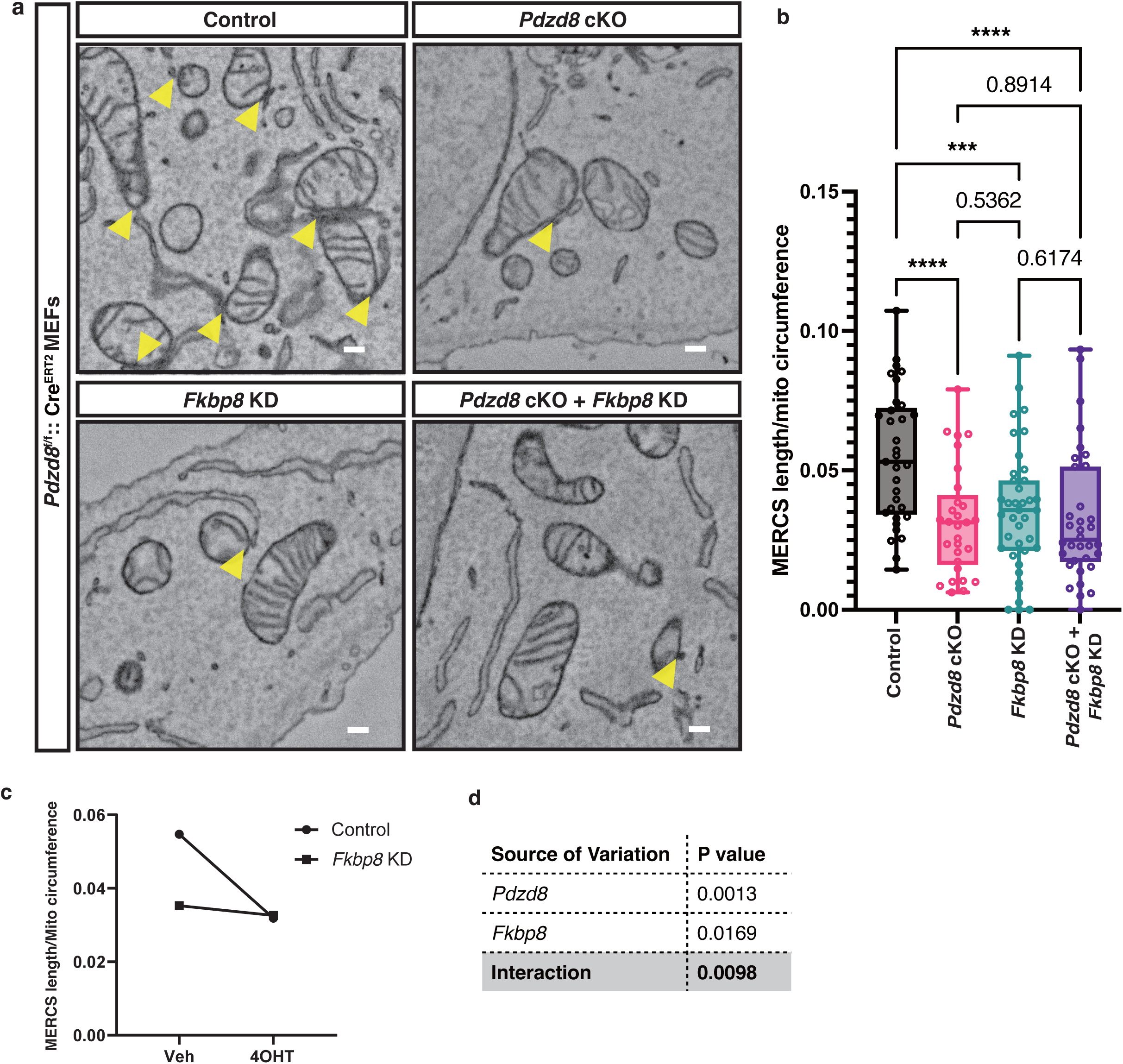
PDZD8 and FKBP8 tether the ER and mitochondria cooperatively. **a,** Representative electron micrographs of *Pdzd8*^f/f^::Cre^ERT2^ MEFs infected with lentivirus carrying shControl or shFKBP8, and treated with or without 0.5 µM 4-OHT. MERCSs (yellow arrowheads) were more frequently observed in the Control cells than in *Pdzd8* cKO, *Fkbp8* KD, and *Pdzd8* cKO + *Fkbp8* KD cells. Scale bar: 200 nm. **b,** Quantification of the MERCS length normalized by the mitochondrial circumference. n = 33, 29, 39, 34 cells from two independent experiments for the control, *Pdzd8* cKO, *Fkbp8* KD, and *Pdzd8* cKO + *Fkbp8* KD cells, respectively. Statistical analysis was performed using One-way ANOVA and Fisher’s LSD test. ****p<0.0001, ***p < 0.001 **c,** The interaction plot corresponding to **c**. Dots show the mean of each condition. **d,** The results of the two-way ANOVA test. The low (< 0.01) variation of the interaction shows that PDZD8 and FKBP8 cooperatively affect the areas of MERCS.

### PDZD8 and FKBP8 colocalize on mitochondria

Although the vast majority of FKBP8 localizes at the mitochondria, an escape of FKBP8 from mitochondria to the ER has been reported upon mitophagy induction^35^. Therefore, to determine if PDZD8 colocalizes with the ER-resident FKBP8 (*cis*-interaction) or with the mitochondrial FKBP8 (*trans*-interaction), we determined the subcellular compartments where PDZD8 and FKBP8 colocalize. Immunostaining using anti-FKBP8 antibodies showed a puncta-like distribution of endogenous FKBP8 and revealed that 88.2% of FKBP8 is localized at mitochondria in HeLa cells (**Fig. 5a, b, Extended Data Fig. 7a**). Calculation of the distance from the centroids of each PDZD8 puncta to the nearest FKBP8 puncta within the mitochondria (‘on mito regions’) revealed that the number of FKBP8 within 450 nm of PDZD8 was considerably higher compared to the control where the centroids of FKBP8 puncta were placed at scrambled positions within ‘on mito regions’. This juxtaposition of PDZD8 and FKBP8 was not observed outside the mitochondria (‘off mito regions’) (**Fig. 5c**). These suggest that FKBP8 localizes near PDZD8 within mitochondria but not in other cytoplasmic region. Additionally, we found that the ratio of PDZD8 intensity overlapped with FKBP8 on mitochondria was significantly reduced by the FKBP8 randomizing (**Fig. 5d**). Moreover, PDZD8 puncta were significantly enriched on the FKBP8-positive area of mitochondria (**Fig. 5e**), indicating that PDZD8 colocalizes with FKBP8 more frequently than random occurrences on mitochondria. To independently confirm these results in cells derived from a different species and using endogenous tagging of FKBP8 and PDZD8 simultaneously, we developed a dual KI strategy (**Extended Data Fig. 7b**), whereby an HA-tag was knocked-in at the *Fkbp8* genomic locus to express HA-FKBP8 in the *Pdzd8*-Venus KI NIH3T3 cell line. Consistent with the localization in HeLa cells, PDZD8 significantly accumulated to FKBP8-present regions on mitochondria compared to FKBP8-absent regions (**Extended Data Fig. 7c-e**). Taken together, these results strongly suggest that an ER-resident protein PDZD8 colocalizes with FKBP8 specifically on the mitochondria.

**Figure 5.**
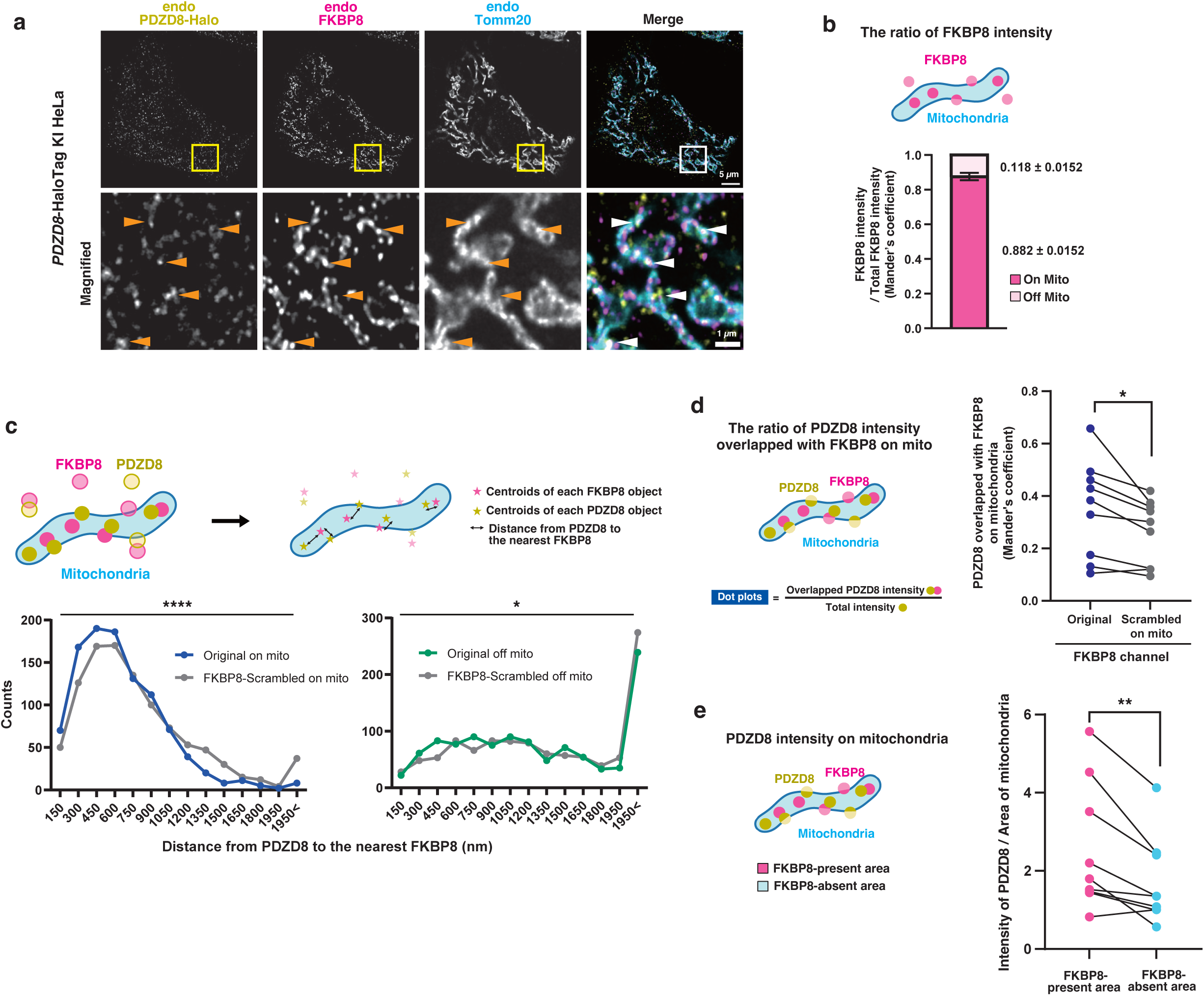
Endogenous PDZD8 and FKBP8 colocalize on mitochondria. **a,** Immunofluorescence analysis of *PDZD8*-Halotag KI HeLa cells. The cells were treated with 200 nM of Janelia Fluor 549 for 20 hours and then stained with antibodies to FKBP8 and to Tomm20. The boxed regions of the top panels are shown at higher magnification in the corresponding lower panels. Arrowheads indicate PDZD8 colocalized both with FKBP8 and Tomm20. Scale bars, 5 μm (original) or 1 μm (magnified). **b,** The ratios of FKBP8 intensity on or outside (off) the mitochondria were determined for images obtained as described in **a**. Error bar is mean ± s.e.m. of nine cells from two independent experiments. The average of three cytoplasmic regions cropped from each of the nine cells were used for the analysis. **c,** Distribution of PDZD8 puncta with indicated distance to the nearest FKBP8 puncta was determined for images obtained as described in **a**. The distance from centroids of each PDZD8 puncta to the nearest FKBP8 centroids was calculated within mitochondria (on mito) or outside of the mitochondria (off mito) respectively. The scrambled FKBP8 centroids were created by shuffling pixels within mitochondria or outside of the mitochondria in the images describing FKBP8 centroids. 9 cells from two independent experiments were used in the calculation. Kolmogorov–Smirnov test was used to test statistical significance. *\*\*\*\*P* < 0.0001, *\*P* < 0.05. **d,** The ratios of PDZD8 intensity overlapped with FKBP8 on mitochondria (Mander’s coefficients) were determined for images as described in **a**. The scrambled FKBP8 images were created by shuffling pixels within mitochondria in the FKBP8 channel. Data are representative of two independent experiments (9 cells). Paired t-test was used to test statistical significance. \**P* < 0.05. **e,** The means of PDZD8 intensity in the FKBP8-present or FKBP8-absent area on mitochondria were determined for images as in **a**. Data are representative of two independent experiments (9 cells). Paired t-test was used to test statistical significance. \*\**P* < 0.01.

### Overexpression of mitochondrial FKBP8 recruits PDZD8 in the proximity of mitochondria

We next tested if overexpression of the mitochondrial FKBP8 is sufficient for recruiting endogenous ER-localized PDZD8 to mitochondria. We overexpressed a mutated form of FKBP8 previously shown to lock its localization at the OMM (FKBP8^N403K^)^35^ in the *PDZD8*-Halotag KI HeLa cells. Strikingly, the overlap of PDZD8 with an OMM-marker YFP-ActA in HA-FKBP8^N403K^ overexpressing cells was significantly increased (**Fig. 6a, b**). This suggests that the mitochondrial FKBP8 binds to PDZD8. Then, we examined if overexpression of FKBP8^N403K^ recruits the ER together with PDZD8 by a correlative light-electron microscopy (CLEM) analysis (**Extended Data Fig. 8a**). Endogenous PDZD8 was labeled with JF549 dye in the *PDZD8*-HaloTag KI HeLa cell expressing with Venus-FKBP8^N403K^ or YFP-ActA (OMM marker). Confocal microscopy with a NSPARC detector was used to visualize JF549 labeled *PDZD8*-HaloTag and Venus-FKBP8^N403K^ or YFP-ActA signals within fixed cells, and the area imaged by confocal microscopy was subsequently re-identified in EM images (**Extended Data Fig. 8b-e**). OMM and the ER membrane within 25 nm of each other (MERCS) were segmented in the EM images and then 3D-reconstructed from 8 slices with 50 nm thickness (total 400 nm thick in z axis) (**Fig. 6c, Extended Data Fig. 8d, e**). The 3D-reconstructed mitochondria and MERCS was aligned to the confocal microscopy image using the FKBP8 signals or ActA signals as landmarks for mitochondria (**Fig. 6d, Supplementary video 4)**. Notably, PDZD8 puncta observed near mitochondria were highly accumulated in MERCS of the FKBP8^N403K^-overexpressing cell (arrowheads in **Fig. 6e, Supplementary video 4**). Taken together, using multiple independent approaches, our results demonstrate that PDZD8 and FKBP8 form a complex between the ER and mitochondria and the overexpression of FKBP8 at OMM increases the abundance of this protein complex at MERCS.

**Figure 6.**
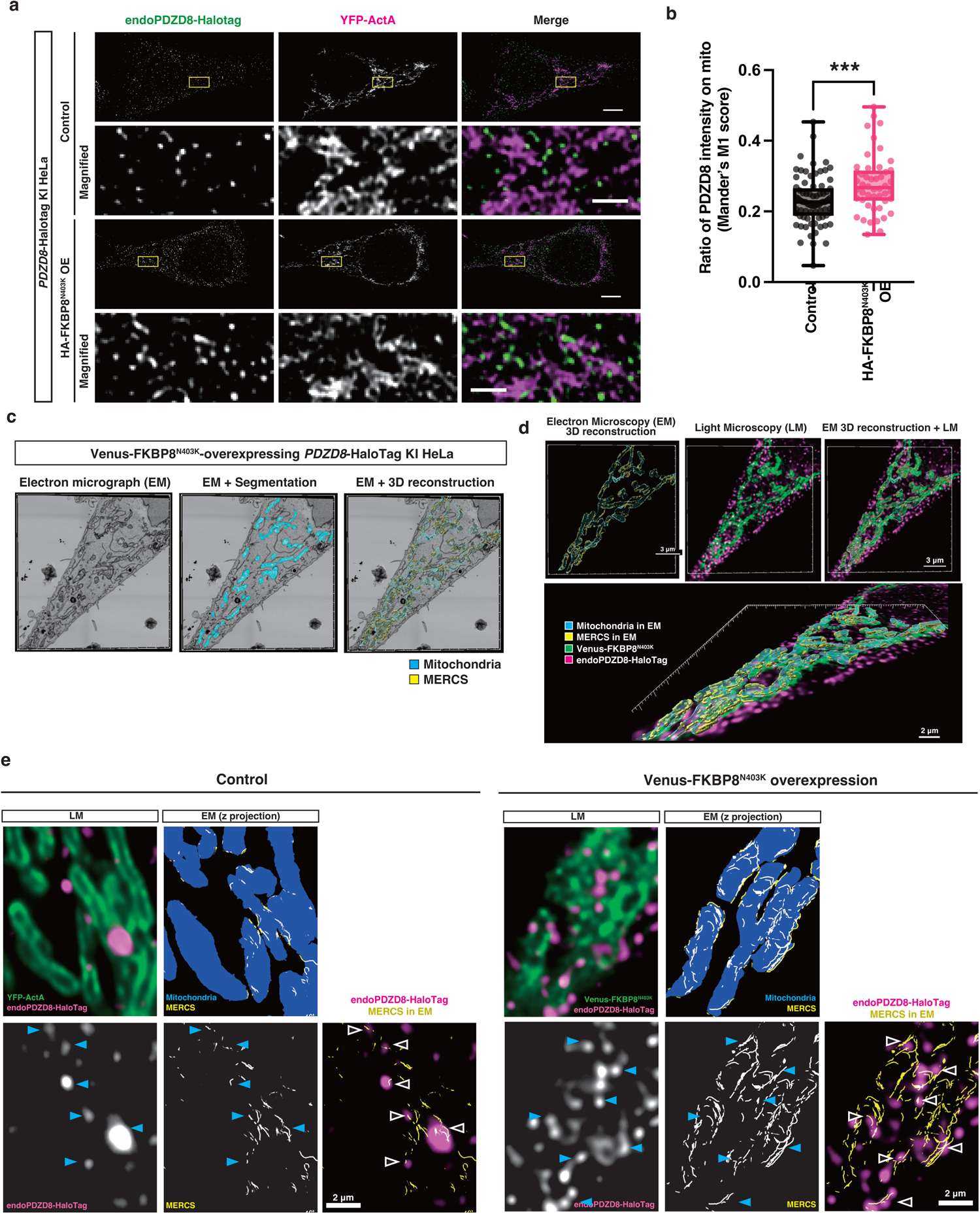
Overexpression of the mitochondrial FKBP8 recruits endogenous PDZD8 to the mitochondrial proximity. **a,** Immunofluorescence analysis of *PDZD8*-Halotag KI HeLa cells overexpressing HA-FKBP8^N403K^. Cells transfected with either the control plasmid (upper two rows) or the plasmid encoding HA-FKBP8^N403K^ (bottom two rows), along with the mitochondrial marker YFP-ActA, were treated with 200 nM of JF549 for 20 hours, and subsequently, the fixed cells were observed using a confocal microscope equipped with a Nikon Spatial Array Confocal (NSPARC) detector. Scale bar: 5 µm (upper panels), 1 µm (lower magnified panels). **b,** Quantification of the percentage of endogenous PDZD8-Halotag intensity overlapping with mitochondria (YFP-ActA-positive area). n = 78, 48 cells for the control and FKBP8 KO cells from two independent experiments. Statistical analysis was performed using Student’s t-test. \*\*\**P* < 0.001 **c–e,** Correlative light and electron microscopy (CLEM) analysis in a *PDZD8*-HaloTag KI HeLa cell. Cells overexpressing with Venus-FKBP8^N403K^ or YFP-ActA (for the control) were treated with 200 nM of JF549 for 20 hours and then fixed cells were observed by a confocal microscope. After that, ultra-thin sections (50 nm thick) were created and observed in a field emission scanning electron microscope (FE-SEM). Electron micrographs of the serial 8 slices were corresponding to an optical section of fluorescence images. Segmentations and 3-demensional (3D) reconstructions of mitochondria and the ER within 25 nm of mitochondria (MERCS) in electron micrographs were shown in **c**. 3D reconstruction from electron micrographs (shown as “EM”) were merged with fluorescence images (shown as “LM”) in **d**. The z projection of mitochondria and MERCS in EM was overlayed with fluorescence images in **e**. Arrowheads indicate PDZD8 puncta that localize to MERCS.

### Overexpression of mitochondrial FKBP8^N403K^ narrows the distance between ER and mitochondria at MERCS

Cryo-electron tomography (cryo-ET) provides the resolution range of 3-50 Å that is not accessible with other techniques and, importantly, allows the quantification of ultrastructural features of MERCS in situ, in native, unfixed conditions. Using correlative cryo-light microscopy and cryo-ET, we studied the *in situ* topology of mammalian MERCS. First, we overexpressed FKBP8^N403K^ in *Pdzd8*-Venus KI NIH3T3 cells and replicated the effect of recruiting and stabilizing PDZD8 in cells grown on cryo-EM grids (**Fig. 7a**). We then used cryo-focused ion beam (cryo-FIB) milling to generate lamellae (<200 nm-thick slice per cell) from *PDZD8*-HaloTag KI HeLa cells overexpressing mScarlet-FKBP8^N403K^ (**Fig. 7b–e**). Cryo-fluorescence images of the lamellae overlaid on their corresponding high-resolution medium-mag TEM montage confirmed the presence of FKBP8^N403K^-mScarlet on all mitochondria including MERCS. Since mScarlet-FKBP8^N403K^ overexpression significantly increased the number of MERCS in each lamella compared to control cells, we fully segmented and labelled membrane structures at 20 MERCS in the FKBP8^N403K^ overexpression condition, and 6 MERCS in the control cells. Using surface morphometrics analysis^36^, we quantified ER-OMM distances at the level of a fraction of MERCS. Our cryo-ET analyses demonstrate that ER-OMM distances at any given MERCS is quite heterogeneous ranging from 10 nm to 50 nm (**Fig. 7f**, Extended Data Fig. 9). Moreover, an aggregate analysis for both the overexpressed and control conditions showed that overexpression of FKBP8^N403K^ significantly (p<0.0005, Kolmogorov-Smirnov test) shifted ER-OMM distances to shorter values with a weighted median value at 25.7 nm compared to 30.1 nm in control MERCS (**Fig. 7g**). These results suggest that overexpression of FKBP8^N403K^, which efficiently recruits ER-localized PDZD8 to MERCS, imposes distances shorter than 25 nm between the ER and OMM membranes at MERCS.

**Figure 7.**
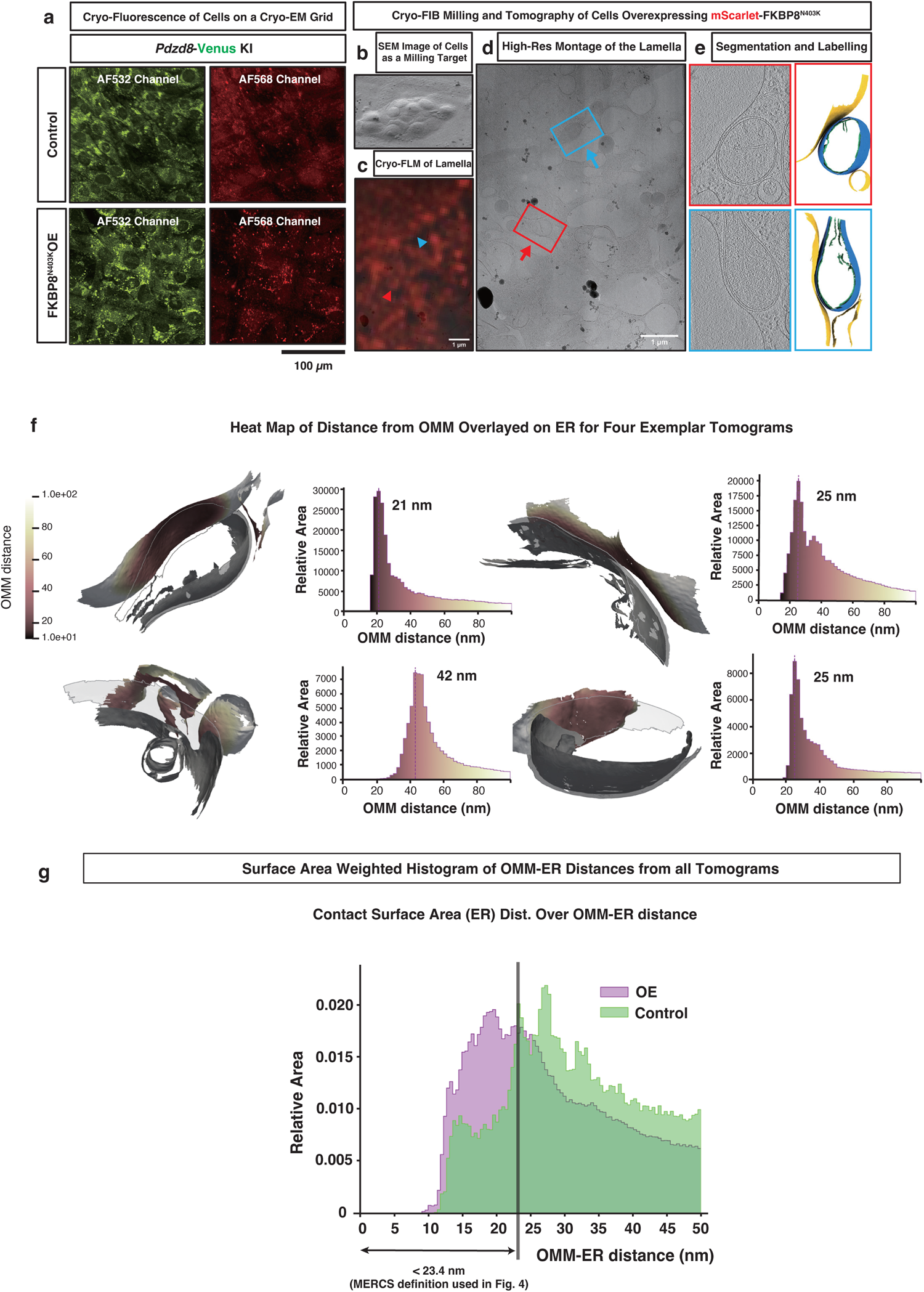
Overexpression of the mitochondrial FKBP8 narrows the ER-OMM distance at MERCS. **a,** Overexpression of FKBP8^N403K^ increases the intensity and the abundance of the PDZD8-Venus puncta. In cryogenic temperatures, autofluorescence of the NIH3T3 cells in the green channel is strong, hence the presence of puncta in the control cells. The Venus fluorophore also emits light in the red channel. **b-e,** For the cryo-ET analysis, mScarlet-FKBP8^N403K^ overexpression (OE) was used to increase the number of MERCS captured in cryo-FIB milled lamellae. An SEM image of a target cell before Cryo-FIB milling is shown (**b**). Cryo-Fluorescence imaging of lamellae confirmed the presence of mScarlet-FKBP8^N403K^ on all the mitochondria (**c**). Using medium-mag high-resolution TEM montages of the lamellae, MERCS were targeted for high-resolution tilt series acquisition (**d**). 80 tomograms containing MERCS were obtained for the OE condition (of which 20 were fully segmented and labelled), and 10 tomograms containing MERCS were obtained for the control (of which 6 were fully segmented and labelled). Two representative tomograms from the OE condition corresponding to the arrows in panel D are shown (**e**). **f,** Surface morphometrics analysis was used to calculate the ER-OMM distance at MERCS. The distances are shown as a heatmap. Mammalian MERCS show a great deal of heterogeneity in their membrane ultra-structure. **g,** Aggregate analysis of the area-weighted ER-OMM distance histogram shows a shift to smaller distances in the overexpression condition compared to the control.

### PDZD8 enhances mitochondrial complexity by suppressing FKBP8

MERCS represent hotspots for both fission and fusion of mitochondria^37, 38^ ^39^. Importantly, previous reports suggested that FKBP8 promotes mitochondrial fission^40, 41^. Thus, we decided to examine the role of PDZD8-FKBP8 complex in the regulation of mitochondrial morphology. Quantitative volume analyses of mitochondria reconstructed from serial electron microscopy images revealed that mitochondria in PDZD8 cKO cells were significantly spherical represented by the smaller mitochondria complexity index (MCI^42^) compared to the control (**Fig. 8a–c**, **Extended Data Fig. 10**). In contrast, mitochondria in FKBP8 KD cells were more elongated and branched compared to the control, resulting in higher MCI (**Fig. 8b, c**). These data suggest that PDZD8 increases but FKBP8 decreases the mitochondrial complexity. Interestingly, PDZD8 KO did not reduce the MCI in FKBP8 KD background (**Fig. 8b, c**). This indicate that PDZD8 suppresses the function of FKBP8 in regulating mitochondrial structure.

**Figure 8.**
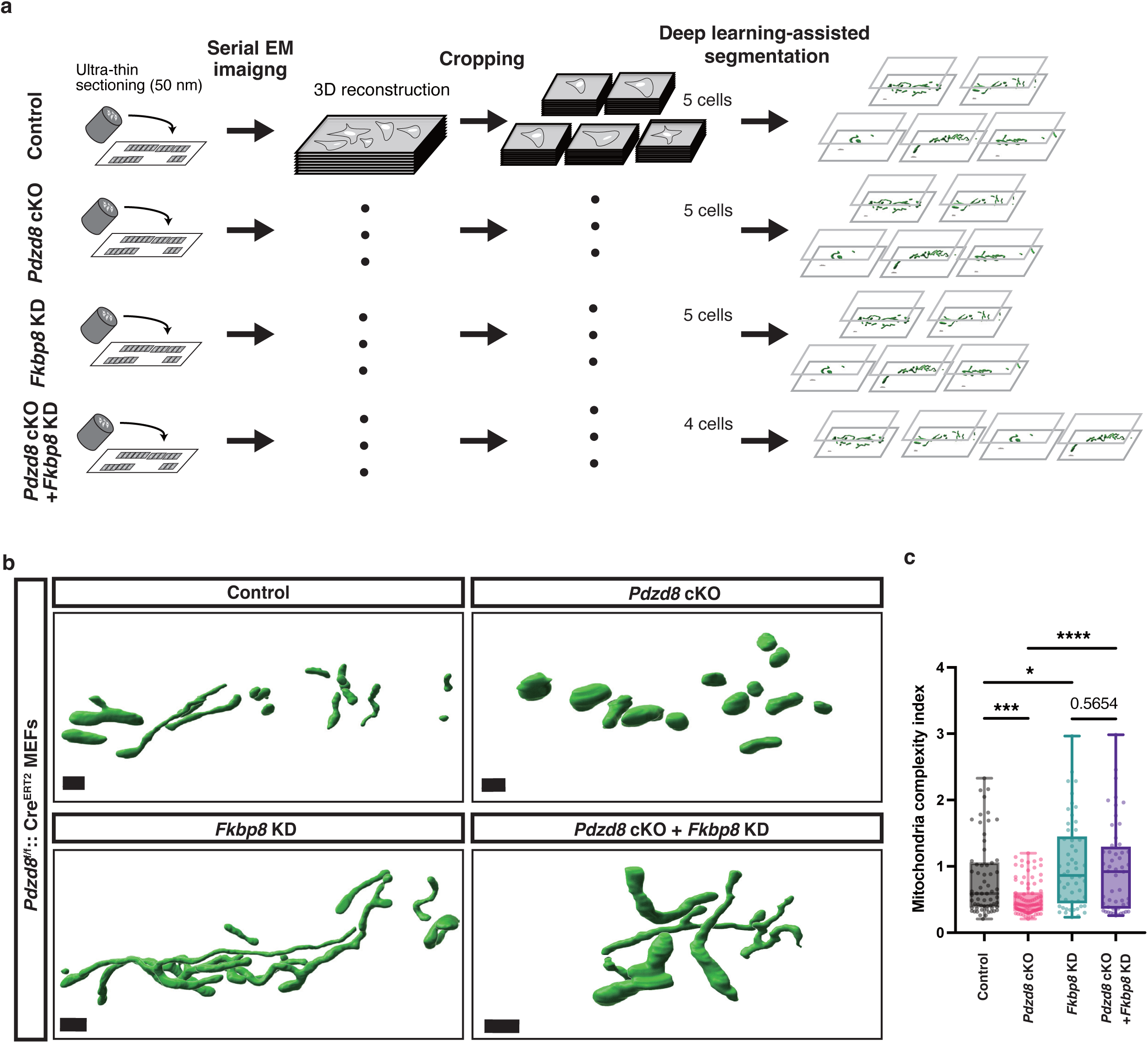
PDZD8 promotes mitochondrial complexity by inhibiting FKBP8. **a,** Schematic of mitochondrial morphology analysis by the volume EM. Serial electron micrographs were obtained by imaging the serial sections with an FE-SEM. After cropping the volume, the mitochondria in the volume were semi-automatically extracted with an AI-assisted pipeline^42^. **b,** Representative 3D reconstruction of mitochondria extracted from serial EM images acquired by array tomography in *Pdzd8*^f/f^::Cre^ERT2^ MEFs infected with lentivirus carrying shControl or shFKBP8, and treated with or without 0.5 µM 4-OHT. Scale bar: 1 µm. **c,** Quantification of MCI (mitochondrial complexity index)^42^. n = 63, 90, 54, and 52 mitochondria from 5, 5, 5, and 4 cells for the control, *Pdzd8* cKO, *Fkbp8* KD, and *Pdzd8* cKO + *Fkbp8* KD cells, respectively. Statistical analysis was performed using One-way ANOVA and Fisher’s LSD test. ****p<0.0001, ***p<0.001, *p < 0.05

## Discussion

The biology of organelle contacts has emerged as molecularly complex and highly dynamic but mediating many crucial aspects of cell physiology. In particular, MERCS play key roles in exchanging Ca^2+^ and lipid between the ER and mitochondria,^8^ regulating mitochondrial fission/fusion, and for biogenesis of autophagosomes ^1, 3, 43, 44^. In this study, we investigated the molecular mechanisms of MERCS formation and its role in regulating mitochondrial morphology by analyzing the dynamics of the ER-mitochondria tethering protein PDZD8 using endogenous tagging, single particle tracking, identifying the partner protein on the mitochondrial side, and investigating mitochondrial morphology at nanometer scale.

### MERCS formation by the PDZD8-FKBP8 protein complex

Our results show (1) specific PDZD8 recruitment at MERCS under endogenous conditions, (2) PDZD8 moves dynamically along the ER membrane and exhibits a significant increase in dwell time or transient ‘capture’ near points of contacts with mitochondria, and (3) using a battery of endogenous tags and biotin ligase-mediated proteomics that FKBP8 is a novel PDZD8 binding partner mediating its tethering function at MERCS. Super-resolution optical imaging and CLEM analysis revealed that FKBP8 is necessary and sufficient for recruiting PDZD8 to MERCS. Furthermore, our ultrastructural analysis suggested that the binding between FKBP8 and PDZD8 is necessary for the formation of a significant fraction of MERCS. We took advantage of a cryo-ET pipeline for characterization of mammalian MERCS at sub-nanometer resolution in native, unfixed, conditions and discovered that the ER-OMM distances are highly variable with a range of 10–50 nm and narrowed by overexpressing FKBP8^N403K^. Taken together, our results revealed a novel molecular mechanism underlying the formation of contacts between the ER and mitochondria.

### Dynamics and localization of PDZD8

The ER spreads throughout the cell and works as a hub exchanging a wide variety of molecules with other organelles especially through membrane contact sites. It has been shown that PDZD8 is an ER protein required for the formation of MERCS, but also localizes at MCS between ER-lysosome, ER-late endosome, and at the ER-late endosome-mitochondria tripartite contacts, all of which we can observe with some frequency in our data^11, 20, 22^. The results of sptPALM showing the dynamic exchange of PDZD8 inside and outside hotspots at ER-mitochondrial contacts suggest that PDZD8 may be able to move rapidly between different types of MCS, as suggested for VAPB. One mechanism that could help facilitate recruitment of PDZD8 to the late endosome is its ability to directly bind to Rab7. Of note, the relatively low binding affinity of FKBP8-PDZD8 *in vitro* revealed in this study, is of the same order of magnitude as that reported for Rab7-PDZD8 *in vitro*^34^. Thus, it is possible that FKBP8 and Rab7 are competing for sequestration of PDZD8 and therefore might control the balance between the areas of MERCS and ER-Lysosome contacts.

### Potential roles of the PDZD8 complex formation in regulating FKBP8 functions

Previous studies revealed that FKBP8 expression induces mitochondrial fragmentation and mitophagy through its LC3-interacting region motif-like sequence (LIRL) and LC3-interacting region (LIR), respectively^40^. In agreement with this, our high-throughput volume EM analysis demonstrated that FKBP8 limits mitochondrial volume maybe by limited fusion or promoting fission (**Extended Data Fig. 10a**). Together with the fact that MERCS provides isolation membranes for autophagy^45^, the MERCS formation via the PDZD8-FKBP8 complex might be a prerequisite for the onset of mitophagy. Of note, a combination of FKBP8 KD and PDZD8 KO revealed that PDZD8 enhances mitochondrial complexity through inhibition of FKBP8. Considering that PDZD8 is required to suppress mitophagy in Drosophila neurons^12^, one potential function of PDZD8 binding to FKBP8 could be to arrest the progression of mitophagy by inhibiting FKBP8-dependent modulation of mitochondrial shape after initiating the formation of the isolation membrane. The CLEM analysis in this study showed that the overexpression of the mitochondrial FKBP8 brought PDZD8 on the positive curvature of mitochondria (**Extended Data Fig. 8f**). Future studies using high resolution live imaging will be required to clarify how the localization of this complex affects the morphological changes of mitochondria.

This study elucidates a molecular pathway that regulates mitochondrial morphology at the interface with the ER. Given the dynamic nature of MERCS, revealing how cellular conditions utilize this pathway to modulate mitochondria will provide novel insights into cellular homeostasis.

## Acknowledgment

We thank Drs. Heike Blockus, Tommy Lewis, Seok-Kyu Kwon for their critical reading of the manuscript and members of the Hirabayashi lab for constructive discussions. We thank Chenxing Jiang and Machiko Tsumura for their technical support. We thank Drs. Yoshibumi Yamaguchi (Hokkaido University), Masato Ohtsuka (Tokai University), Masayuki Miura (The University of Tokyo), and Makoto Matsuyama (Shigei Medical Research Institute) for the kind instruction of the iGONAD method, Dr. Luke Hammond (Columbia University) for the kind instruction of fluorescence image analysis, and Drs. Satoru Takahashi, Yoko Ishida, Chieko Saito, Ikuko Koyama-Honda, and Noboru Mizushima (The University of Tokyo) for the kind instruction of the CLEM method. We thank Drs. Shigeo Okabe and Yuka Sato for the support of FE-SEM imaging. We thank Dr. Jonathon Nixon-Abell for performing pilot experiments with sptPALM of PDZD8 and assisting with establishing imaging and tracking conditions. We acknowledge Shunsuke Kihara for the support of NIKON NSPARC microcopy. This work was supported by JSPS KAKENHI under Grant Number JP20H04898 (Y.H.), JP22H05532 (Y.H.), JP21J00490 (S. A-I.), JP22J23099 (K.N.), AMED under Grant number JP19dm0207082 (Y.H.), JP21wm0525015 (Y.H.), SECOM Science and Technology Foundation Research grant (Y.H.), the Uehara memorial foundation research grant (Y.H.), the Naito Foundation (S. A-I.), NIH-NINDS R35 award NS127232 (F.P.), Joint Usage and Joint Research Programs by the Institute of Advanced Medical Sciences of Tokushima University (K.N., T.N., Y.S-S., Y.H., and H.K.), and The University of Tokyo WINGS-LST “Collaboration project” and “Laboratory Practice” (K.N. and Y.H.). We thank Dr. Chrisostomos Prodromou (University of Sussex) for providing the pRSETA-hFKBP8 (1-380) - Histag. We thank Drs. Yuji Tsunekawa and Fumio Matsuzaki (RIKEN) for providing the YT210 plasmid. pENTR4-HaloTag (w876-1) was a gift from Dr. Eric Campeau (Addgene plasmid #29644; http://n2t.net/addgene:29644; RRID: Addgene_29644).

## Author contribution

Y.H. conceptualized and supervised the project. K.N., S. A-I., T.N., Y.S-S., Y.D., S.S., M.T., and Y.H. designed and performed the experiments. M.P. performed cryo-ET analysis advised by C.P., B.C., and F.P.. J.J. helped with the cryo-ET grid preparation and data processing. C.J.O. and J.L.-S. performed sptPALM analysis. K.N. performed *in vitro* analysis under the supervision of M.N. and K.T.. Y.K. and Y.G. generated *Pdzd8*-3×HA KI mice. H.K. acquired and analyzed the MS data. C.K., H-W. R, J.K.S. performed proximity labeling-MS experiment of *Pdzd8*-TurboID KI Neuro2a. K.N., S. A-I., T.N., and Y.H. wrote the manuscript with the help of the rest of the authors.

## Methods

### Cell culture and plasmid transfection

NIH3T3 cells, Neuro2a cells, HeLa cells, and MEFs were maintained with Dulbecco’s Modified Eagle Medium (DMEM, Sigma-Aldrich, catalog no. D6429) supplemented with 10% FBS (MP Biomedicals, catalog no. 2917346), and 1% Penicillin-Streptomycin (Gibco, catalog no. 15140-122) at 37°C under 5% CO_2_. COS7 cells were maintained in phenol red-free DMEM (Corning, catalog no. 25200114) supplemented with 10 % FBS (Corning, catalog no. 35-011-CV), 1% Penicillin-Streptomycin, and 1% L-glutamine (Corning, catalog no. 25-005-CI). NIH3T3 cells, Neuro2a cells, and MEFs were transfected with plasmids by polyethyleneimine (Polysciences). HeLa cells were transfected with plasmids by Lipofectamine LTX reagent with Plus reagent (Thermo Fisher) and Lipofectamine 2000 transfection reagent (Thermo Fisher), or with siRNAs by Lipofectamine RNAiMAX transfection reagent (Thermo Fisher). All DNA plasmids used in this work are listed in Supplemental Table 4.

### Animals

All animals were maintained and studied according to protocols approved by the Animal Care and Use Committee of The University of Tokyo.

### Generation of Pdzd8-Venus/TurboID/Halotag knock-in cell lines

*Pdzd8*-Venus KI NIH3T3, *Pdzd8*-TurboID KI Neuro2a cells, *PDZD8*-TurboID KI HeLa cells, and *PDZD8*-Halotag KI HeLa cells are generated as previously described.^8^ The plasmids pCAG-mPDZD8_cterm-Venus-P2A-Neo for *Pdzd8*-Venus KI NIH3T3 cells, pCAG-mPDZD8_cterm-v5-TurboID-P2A-Neo for *Pdzd8*-TurboID KI Neuro2a cells, pCAG-hPDZD8_cterm-v5-TurboID-P2A-Neo for *PDZD8*-TurboID KI HeLa cells, and pCAG-hPDZD8_cterm-HaloTag-P2A-Neo for *PDZD8*-HaloTag KI HeLa cells were used as donor vectors respectively.

### Generation of Pdzd8-Venus and HA-Fkbp8 double knock-in cell lines

For CRISPR-Cas9 plasmid, CRISPR guide RNA that targets the region prior to *Fkbp8* start codon was designed using CRISPR Design tool (Horizon Discovery Ltd.) and cloned into pSpCas9(BB)-2A-Puro (PX459) V2.0 (Addgene plasmid # 62988) as previously described.^46^ The donor oligonucleotides containing the 5’ arm sequence, the sequence of HA tag and the 3’ arm sequence (5’ - TCC CCG AGC CGC AGG GCC AGT TCC TGA TCC CAG CAG CAT GTA CCC ATA CGA TGT TCC AGA TTA CGC TGC GTC TTG GGC TGA GCCCTC TGA GCC TGC TGC CCT - 3’) were obtained from Eurofins Genomics. *Pdzd8*-Venus KI NIH3T3 cells were transfected with CRISPR-Cas9 plasmid and the donor oligonucleotides by polyethylenimine.

### Generation of tamoxifen-inducible Pdzd8 conditional KO cell lines (Pdzd8^f/f^::Cre^ERT2^ MEFs)

Mouse embryos were dissected from anesthetized females at embryonic day 13.5. The embryos were minced, and after treatment with 0.25% trypsin (Gibco), 50 μg/mL of DNaseI (Merck) and 0.67 mg/mL of Hyaluronidase (Merck) in PBS for 20 minutes, the cells of the resulting suspension were plated onto 100-mm culture dishes and maintained in culture medium for 4 days. The cells were then immortalized by transfecting with plasmids encoding simian virus 40 (SV40) large T antigen (pMK16_SV40 T ori (-)^47^). After that, the cells were infected with lentivirus carrying Cre-ERT2 and single cell clones were obtained using a limiting dilution in 96-well plates.

### Generation of Pdzd8-3×HA knock-in mice

CRISPR/Cas9-mediated genome editing was performed using the iGONAD method as previously reported.^48^ Briefly, 2- to 3-month old female mice (Jcl:ICR, CLEA Japan) were mated with male mice the day before electroporation. The female mice with virginal plug were used for iGONAD at embryonic day 0.75. Genome editing solution was prepared with 1 mg/ml Cas9 protein (IDT, 1081059), 30 mM crRNA (annealed with tracrRNA, IDT, 1072534), 2 mg/ml ssODN (IDT, Ultramer DNA Oligo, standard desalting), and FastGreen (Fujifilm Wako, 061-00031) in OPTI-MEM (Thermo Fisher Scientific, 11058021). The oviducts of the female mice were exposed and injected with the solution through microcapillary injection. Oviduct electroporation was performed using NEPA21 and CUY652P2.5×4 (NEPA gene) with the following protocol: three poring pulses (50 V, 5 msec, 50 msec interval, and 10% decay [±pulse orientation]) and three transfer pulses (10 V, 50 msec, 50 msec interval, and 40% decay [±pulse orientation]). After electroporation, the oviducts were returned to their original position. The sequences of crRNA and ssODN were as follows: crRNA, 5’ - ATT GAT TAC ACT GAC TCA GA - 3’ and ssODN, 5’ - AGC CAT TCA GCA ACA TTT CCG ATG ACT TGT TCG GCC CAT CTG AGT CAG TGT ACC CAT ACG ATG TTC CAG ATT ACG CTG GCT ATC CCT ATG ACG TCC CGG ACT ATG CAG GAT CCT ATC CAT ATG ACG TTC CAG ATT ACG CTG TTT AAT CAA TAA GCT ATT TCA ACT TTC ACA TGG ATG GAG GGG ACA AGA CGT A - 3’.

### Abs

Primary Abs for immunostaining; anti-Tomm20 (Abcam, ab78547; 1:500), anti-LAMP1 (BD Bioscience, 553792; 1:500), rat anti-GFP (Nacalai, 04404-84; 1:200-1:500), mouse anti-GFP (Invitrogen, A-11120; 1:1,000), anti-Rab7 (Cell Signaling Technology, 9367; 1:100), anti-OXPHOS complex (Invitrogen, 45-8099; 1:200), anti-FKBP8 (R and D systems, MAB3580; 1:500), anti-HA-tag (Cell Signaling Technology, 3724; 1:200), anti-HA-tag (BioLegend, 16B12; 1:500).

Primary Abs for immunoblotting; anti-PDZD8 (Hirabayashi et al.^8^; 1:500), anti-HA-tag (BioLegend, 901501; 1:2,000), anti-FKBP8 (R and D systems, MAB3580; 1:500), anti-VAPA (Bethyl laboratories, A304-366A; 1:1,000), anti-Mfn2 (Abcam, ab56889; 1:1,000), anti-β-actin (Cell Signaling Technology, 4967; 1:500), anti-FLAG (M2) (Sigma-Aldrich, F1804; 1:1,000), anti-α-tubulin (Sigma-Aldrich, T6188; 1:1,000), anti-GFP (Medical & Biological Laboratories, 598; 1:1,000), anti-His-tag (Medical & Biological Laboratories, D291-3S; 1:1,000), and anti-v5 (Abcam, ab27671; 1:500).

### Immunoblotting

Cells were lysed with a solution containing 20 mM Hepes-NaOH (pH 7.5), 150 mM NaCl, 0.25 M sucrose, 1 mM EDTA, 0.1% SDS, 0.5% Sodium deoxycholate, 0.5% NP-40, 1 mM Na_3_VO_4_, cOmplete Mini Protease Inhibitor Cocktail (Roche) and Benzonase (25 U/ml). Insoluble pellets and supernatants were separated by centrifugation at 15,000 × g at 4°C for 15 minutes. The supernatants were boiled with 1 × Laemmli’s sample buffer containing 10% mercaptoethanol at 98°C for 5 minutes. The cell lysates were fractionated by SDS-PAGE on a 10% gel or a 4-15% gradient gel (Bio-rad) and the separated proteins were transferred to a polyvinylidene difluoride membrane (Merck). The membrane was incubated first with primary Abs for 24 hours at 4°C and then with HRP–conjugated secondary Abs (GE Healthcare) for 1 hour at room temperature. After a wash with TBS-T (50 mM Tris-HCl (pH 8), 150 mM NaCl, and 0.05% Tween 20), the membrane was processed for detection of peroxidase activity with chemiluminescence reagents (100 mM Tris-HCl (pH 8.5), 1.25 mM Luminol, 0.2 mM P-Coumaric Acid, 0.01 % H_2_O_2_) and the signals were detected by Image Quant LAS4000 instrument (GE Healthcare).

### Coimmunoprecipitation analysis

#### Cross-linked IP in *Pdzd8*-3×HA knock-in mouse brain

*Pdzd8*-3×HA knock-in mice and control littermates at postnatal 10 days were put to sleep using medetomidine hydrochloride (Domitor, Nippon zenyaku kogyo, 0.75 mg/kg), midazolam (Sandoz, 4 mg/kg) and butorphanol (Vetorphale, Meiji Seika Pharma Co., Ltd, 5 mg/kg). Pups were then put on the ice for 5 minutes and exsanguinated by terminal intracardial perfusion with ice-cold 2% paraformaldehyde (Merck) in phosphate-buffered saline (PBS). The neocortex was then removed and sonicated five times for 30 seconds with ice-cold lysis buffer (50 mM Tris-HCl (pH 7.5), 1 mM EDTA, 0.2 % Triton-X100, PhosSTOP phosphatase inhibitor (Roche) and cOmplete protease inhibitor cocktail (Roche)). Insoluble pellets and supernatants were separated by centrifugation at 15000 × g at 4°C for 15 minutes. The supernatants were incubated in rotation at 4°C for 20 hours with a protein complex of anti-HA antibody (Cell signaling technology, C29F4) and Sera-Mag SpeedBeads Protein A/G (Cytiva). After the rotation, beads were washed three times with TBS buffer. The immunoprecipitates were eluted from beads by incubating in 2 × Laemmli’s sample buffer containing 10% mercaptoethanol at 98°C for 10 minutes and then subjected to immunoblotting. Total fraction samples were prepared using 2% of the cell extracts.

#### Cross-linked IP in *Pdzd8-*Venus knock-in NIH3T3 cell

Cells were fixed with 0.1% PFA for 10 minutes at room temperature, and 100 mM glycine-NaOH was treated for 4 minutes at RT. Cells were washed twice with ice-cold PBS and lysed with ice-cold lysis buffer (50 mM Tris-HCl, pH 7.4, with 150 mM NaCl, 1 mM EDTA, 0.2% Triton-X100, 1 mM Na_3_VO_4_ and cOmplete Mini Protease Inhibitor Cocktail (Roche)). Cell extracts were incubated on ice for 15 minutes, then insoluble pellets and supernatants were separated by centrifugation at 15,000 × g at 4°C for 15 minutes. The supernatants were incubated in rotation at 4°C for 20 hours with a protein complex of anti-GFP antibody (MBL) and Dynabeads Protein A (Thermo Fisher Scientific). After the rotation, beads were washed three times with TBS buffer. The immunoprecipitates were eluted from beads by incubating in 2×Laemmli’s sample buffer, then mercaptoethanol was added at the final concentration of 9%. Samples were boiled at 98°C for 5 minutes and then subjected to immunoblotting. Total fraction samples were prepared using 1.5% of the cell extracts.

#### IP in PDZD8-3×FLAG and HA-FKBP8 overexpressing cell

*Pdzd8*^f/f^::Cre^ERT2^ MEFs were treated with 1 μM 4-OH tamoxifen for 24 hours and then transfected with plasmids encoding 3×FLAG-tagged full length PDZD8/deletion mutants and HA-tagged FKBP8. 24 hours post transfection cells were lysed with ice-cold lysis buffer (50 mM Tris-HCl (pH 7.5), 1 mM EDTA, 0.2 % Triton-X100, 1 mM Na_3_VO_4_ and cOmplete protease inhibitor cocktail (Roche)), and insoluble pellets and supernatants were separated by centrifugation at 15,000 × g at 4°C for 15 minutes. The supernatants were incubated in rotation at 4°C for more than 3 hours with a protein complex of anti-HA antibody (Cell signaling technology) and Dynabeads Protein A (Thermo Fisher Scientific). After the rotation, beads were washed twice with TBS-T buffer and once with TBS buffer. The immunoprecipitates were eluted from beads by incubating in 2×Laemmli’s sample buffer, then mercaptoethanol was added at the final concentration of 9%. Samples were boiled at 98°C for 5 minutes and then subjected to immunoblotting. Total fraction samples were prepared using 20% of the cell extracts.

#### Knocking out of endogenous FKBP8 by CRISPR/Cas9

The pCAX-Cas9 and gRNA backbone vector (YT210) were generously provided by F. Matsuzaki^49^. To enhance knockout (KO) efficiency, three gRNAs targeting different exons were designed for the CRISPR/Cas9-based KO system as previously documented^50^. The gRNA sequences were designed using either Crispor^51^ or the CRISPRdirect^52^. Target sequences were amplified using forward and reverse oligonucleotides through PCR and subsequently cloned into the gRNA backbone vector at the AflII restriction sites^49^.

#### Immunocytochemistry

Cells were fixed with 4% paraformaldehyde for 15 minutes at 37°C, permeabilized with 90% methanol in PBS for 20 minutes under −20°C and incubated for 20 hours (Fig. 1B and D, Fig. 3J, Fig. S1A, Fig. S3G) or 1 hour (Fig. 5A, Fig S3E) in PBS containing 2% FBS and 2% BSA (blocking buffer) at room temperature. They were then exposed at room temperature first for 1 hour to primary Abs in blocking buffer and then for 30 minutes to Alexa Fluor–conjugated secondary Abs (Thermo Fisher Scientific) in blocking buffer. ProLong Gold (Thermo Fisher Scientific) was used as a mounting medium. Images were acquired on a Nikon Ti2 Eclipse microscope with an A1R confocal, a CFI Plan Apochromat Lambda D 100X Oil (NA 1.45), a laser unit (Nikon; LU-N4, 405, 488, 561, and 640 nm), and filters (450/50 nm, 525/50 nm, 595/50 nm, 700/75 nm for 405 nm, 488 nm, 561 nm, 640 nm laser, respectively). All equipment was controlled via Nikon Elements software. Optical sectioning was performed at Nyquist for the longest wavelength. The resulting images were deconvoluted with NIS-elements (Nikon) and processed with NIS-elements (Nikon) or ImageJ (NIH). In Fig. 3J and Fig. S3G, the images were taken as z-stack images (interval; 100nm) and then 3D-deconvoluted with NIS-elements (Nikon) to enhance resolution. One z-slice image was arbitrarily selected in each cell as a representative image and used in the analyses in Fig. 5b-e, and Fig. S7d and e.

#### Analysis of PDZD8 and FKBP8 localization in fluorescent images

All analyses of PDZD8 or FKBP8 localization in fluorescent images were conducted using homemade programs written with Python, as detailed below. All binarized images were created using OpenCV’s threshold function. In Extended Data Fig.1c and d, mitochondrial area, lysosomal area, or late endosomal area were defined by binarizing signals of Tomm20/OXPHOS, Lamp1, or Rab7, respectively, and then the percentages of PDZD8 intensity on mitochondria, lysosome or late endosome were calculated. In Fig.3f–g, the regions of interest (ROI) were defined as areas containing the transfection marker (pCAG-mtagBFP2^53^). The mitochondrial areas were defined by binarizing signals of Tomm20 and then the percentage of PDZD8 intensity on mitochondria (Mander’s coefficient, M1) was calculated. In Fig.5b-e, mitochondrial areas were defined by binarizing signals of Tomm20. In Fig.5b, to calculate Mander’s coefficient between FKBP8 and Tomm20, the sum of FKBP8 intensity on mitochondria divided by total FKBP8 intensity was calculated. In Fig.5c, using OpenCV’s connectedComponentsWithStats function, the puncta of PDZD8 and FKBP8 were segmented in the binarized images of PDZD8 and FKBP8, respectively, and then obtained the centroids of individual puncta. To define the centroids in the “on mito” or “off mito” regions, the images mapping PDZD8 and FKBP8 centroids were masked using the binarized images of Tomm20. To obtain scrambled images of FKBP8, the pixels of the image mapping FKBP8 centroids on mito or off mito regions were shuffled in the corresponding area using the random module of Python. The distance map of FKBP8 was created using OpenCV’s distanceTransform function in the images with FKBP8 centroids “on mito”, shuffled “on mito”, “off mito”, and shuffled “off mito”, respectively. Then the number of PDZD8 puncta with the distance from the nearest FKBP8 corresponding to each bin indicated in x-axis of the graph was calculated. In Fig.5d, to calculate Mander’s coefficient between PDZD8 and FKBP8 on mitochondria, the sum of PDZD8 intensity on FKBP8-present mitochondrial regions divided by total PDZD8 intensity on mitochondria was calculated. The scrambled images of FKBP8 were created by shuffling the pixels of FKBP8 channel within mitochondrial area using the random module of Python. In Fig.5e, FKBP8-present or FKBP8-absent mitochondrial areas were defined as ROIs using binarized images of Tomm20 and FKBP8, and then the sum of PDZD8 intensity at ROIs divided by the area of ROIs was calculated. Image analyses in Fig. S7d and e were performed using the same methods as in Fig.5d and e. In Fig. 6a–b, the ROI were manually defined as the cytoplasmic region using the YFP-ActA signal as a guide. The mitochondrial area was defined as binarized images of YFP-ActA and then the percentage of PDZD8 intensity on mitochondria (Mander’s coefficient, M1) was calculated.

#### Imaging of PDZD8 on mitochondria in FKBP8^N403K^ overexpressing cells

Cells were fixed with 4% paraformaldehyde for 15 minutes at 37°C. The fixed cells were mounted by ProLong Gold (Thermo Fisher Scientific). Images were acquired on a Nikon Ti2 Eclipse microscope with a Nikon AX confocal microscopy with a Nikon Spatial Array Confocal (NSPARC) detector and a CFI Plan Apochromat Lambda D 100X Oil (NA 1.45). The wavelength range of 430–463 nm, 503–545 nm, or 582–618 nm was obtained by excitation with a 405 nm, 488 nm, or 561 nm laser, respectively.

#### Protein expression and purification

For the expression of FLAG-tagged human PDZD8 (1, 28-) -, human *PDZD8* sequences were cloned into the pCAG vector. Recombinant human PDZD8 (1-28) - FLAG was expressed in Expi293 Cells (Thermo Fisher Scientific) using ExpiFectamine 293 Transfection Kit (Thermo Fisher Scientific) according to the manufacturer’s protocol. The cells were cultured for 4 days after transfection at 37°C and 8% CO_2_. The Expi293 cell pellets were homogenized with lysis buffer (25 mM Tris-HCl (pH 8.0) 150 mM NaCl) and centrifuged at 40,000 × g for 30 minutes at 4°C. The supernatant was filtered through an 0.8-μm pore-size filter and subsequently applied to a DDDDK-tagged protein purification gel (MBL) equilibrated with the lysis buffer. After washing with the lysis buffer once, the FLAG-tagged proteins were eluted with 1 M L-arginine-HCl (pH 4.4). The eluted fraction was dialyzed with an SEC buffer (25 mM Tris pH 8.0 containing 300 mM NaCl, and 5 mM DTT). The dialyzed fraction was subjected to size-exclusion chromatography in a HiLoad 16/600 Superdex 200 pg column equilibrated with the SEC buffer in an AKTA system (GE Healthcare). The purified fractions were concentrated using Amicon Ultra-15 (Cut off: 100 kDa) Centrifugal Filter Units (MERCK). For expressing GST-tagged human PDZD8 (1, 28-506) - HA, human *PDZD8* sequences were cloned in pGEX4-T-1 vector (Cytiva) and transformed into *Escherichia coli* BL21 (DE3) cells. After culturing 24 hours at 37°C, the cells are incubated at 28°C until OD_600_ reaches 0.6-1.0. Then 0.5 mM of IPTG was added into the LB medium and incubated at 20°C. 16–20 hours after IPTG induction, the cells are collected by centrifugation (8,000 × g 10 minutes 4°C), frozen by liquid nitrogen, and stored at −30°C. The frozen pellet was mixed with lysis buffer (20 mM Tris pH 8.0, 500 mM NaCl, 5 mM DTT, 10 mM EDTA, and Benzonase diluted at 1:5000) and lysed with an ultrasonic disruptor. The cell lysate was centrifuged (40,000 × *g* for 30 minutes) at 4°C. The supernatant was filtered through an 0.8-μm pore-size filter and subsequently loaded onto a Glutathione Sepharose 4B (Cytiva) equilibrated with a lysis buffer. After washing with (20 mM Tris pH 8.0, 500 mM NaCl, 5 mM DTT, 10 mM EDTA, 1% Triton-X100) once, with (20 mM Tris pH 8.0, 500 mM NaCl, 5 mM DTT) twice, and the GST-tagged protein were eluted with Elute buffer (50 mM Tris-HCl, 10 mM reduced glutathione, pH 8.0). For the expression of His-tagged human FKBP8 (1-380), pRSETA-hFKBP8 (1-380) - Histag (a kind gift from Dr. Chrisostomos Prodromou^54^) was transformed into *Escherichia coli* C43 (DE3) cells. After culturing for 24 hours at 37°C, the cells were incubated at 28°C until the OD_600_ reached 0.6–1.0. Then, 0.5 mM of IPTG was added to the LB medium and incubated at 20°C. 16-20 hours after IPTG induction, the cells were collected by centrifugation (8,000 × g for 10 minutes at 4°C), frozen in liquid nitrogen, and stored at −30°C. The frozen pellet was homogenized with lysis buffer (20 mM Tris-HCl pH 7.5, 100 mM NaCl, 0.5 mM imidazole, Benzonase diluted at 1:10,000) and centrifuged at 40,000 × g for 30 minutes at 4°C. The supernatant was filtered through an 0.8-μm pore-size filter and subsequently loaded onto a TALON Metal Affinity Resin (Clontech) equilibrated with the lysis buffer. After washing with the wash buffer (20 mM Tris-HCl pH 7.5, 100 mM NaCl 5 mM imidazole) twice, the protein was eluted with elution buffer (20 mM Tris-HCl pH 7.0, 100 mM NaCl, 500 mM imidazole). The eluted fraction was dialyzed with an SEC buffer (20 mM Tris pH 7.5 containing 500 mM NaCl, 1.0 mM EDTA, and 5 mM DTT). The dialyzed fraction was subjected to size-exclusion chromatography in a HiLoad 16/600 Superdex 200 pg column equilibrated with the SEC buffer in an AKTA system (GE Healthcare).

#### SPR (Surface Plasmon Resonance)

The interactions of hPDZD8 (1, 28-) - FLAG with hFKBP8 (1-380) - Histag were analyzed using SPR in a Biacore T200 instrument (Cytiva). A Series S CM5 Biacore sensor chip (Cytiva) was activated with N-hydroxysuccinimide/N-ethyl-Ń-(3-dimethylaminopropyl) carbodiimide hydrochloride, followed by immobilization of hPDZD8 (1, 28-) - FLAG at 618 resonance units. After the immobilization, the activated surface of the sensor chip was blocked with 1 M ethanolamine hydrochloride (pH 8.5). Binding analysis was performed at 25°C in a running buffer of HBS-T (10 mM HEPES-NaOH, pH 7.4, 150 mM NaCl, and 0.005% (v/v) Tween-20). A series of five 2.5-fold dilutions of the FKBP8 solution was injected into the sensor chip at 30 μL/min, with a contact time of 120 seconds and a dissociation time of 120 seconds. The K_D_ values were calculated with the Steady State Affinity model on Biacore T200 Evaluation Software, version 3.2 (Cytiva).

#### GST-Pulldown assay

GST-hPDZD8 (1, 28-506)-HA and hFKBP8 (1-380)-Histag were mixed with Glutathione Sepharose 4B (Cytiva) in a buffer (20 mM Tris pH 7.5 containing 500 mM NaCl, 1.0 mM EDTA, 5 mM DTT) and incubated for 3 hours at 4°C. After washing with a wash buffer (20 mM Tris pH 7.5 containing 500 mM NaCl, 5mM DTT) twice, the proteins were eluted by cleaving the thrombin cleavage site with 0.04 U/ µL thrombin (Cytiva) in an elution buffer (20 mM Tris pH 7.5 containing 500 mM NaCl, 5 mM DTT, 2 mM MgCl_2_). The eluate was subjected to SDS-PAGE and processed for Western blotting with anti-Histag and anti-HA antibodies.

#### Electron microscopy

The cells were fixed with 2.5% glutaraldehyde (Electron Microscopy Sciences) in DMEM, for 1 hour at 37°C. After being washed with 0.1 M phosphate buffer (0.02 M sodium dihydrogenphosphate dihydrate, 0.08 M disodium hydrogenphosphate), the cells were scraped and collected with 0.2% BSA/0.1 M phosphate buffer followed by centrifugation at 820 × g. After being embedded in low melting agarose (2% in 0.1 M phosphate buffer, MP Biomedicals), cell pellets were sectioned at 150-µm thickness with a Leica VT1000S vibratome. The sections were post-fixed with 1% OsO_4_ (Electron Microscopy Sciences) and 1.5% potassium ferrocyanide (FUJIFILM Wako Pure Chemical Corporation) in a 0.05 M phosphate buffer for 30 minutes. After being rinsed for 3 times with H_2_O, the cells were stained with 1% thiocarbohydrazide (Sigma-Aldrich) for 5 minutes. After being rinsed with H_2_O for 3 times, cells were stained with 1% OsO_4_ in H_2_O for 30 minutes. After being rinsed with H_2_O for 2 times at room temperature and 3 times with H_2_O at 50°C, the cells were treated with Walton’s lead aspartate (0.635% lead nitrate (Sigma-Aldrich), 0.4% aspartic acid (pH 5.2, Sigma-Aldrich)) at 50°C for 20 minutes. The sections were followed by incubations in an ascending ethanol series (15 minutes each in 50% on ice, 70% on ice, and 10 minutes each in 90%, 95% ethanol/H_2_O at room temperature), 10 minutes in 100% ethanol 4 times and 60 minutes in butyl 2,3-epoxypropyl ether (Fujifilm Wako pure chemical corporation). This was followed by infiltration of Epok812 resin-butyl 2,3-epoxypropyl ether for 24 hours at a 1:1 dilution. After incubating with 100% Epok812 resin for 4 hours, followed by 2 hours, the resin was cured at 40°C for 12 hours, followed by 60°C for 48 hours. Epok812 resin was made by mixing 7.5 g of MNA (Oken), 13.7 g of Epok812 (Oken), 3.8 g of DDSA (Oken), and 0.2 g of DMP-30 (Oken). Resin blocks were trimmed with a TrimTool diamond knife (Trim 45; DiATOME). 50-80 nm thick ultra-thin sections made with a diamond knife (Ultra 45; DiATOME) were collected on a cleaned silicon wafer strip in a Leica Ultramicrotome (UC7). For Fig. 6, the ultra-thin sections were imaged with a scanning electron microscope (JSM7100F; JEOL). Imaging was done at 5 kV accelerating voltage, probe current setting 12, 1,280 × 960 frame size, and 7.4-mm working distance, using the Backscattered Electron Detector. The final pixel size was a 7.8 nm square. In the array tomography analysis, resin blocks were serially sectioned at a 50-nm thickness with a diamond knife (Ultra JUMBO, 45 degrees; DiATOME) fitted in a Leica Ultramicrotome (UC7) to obtain a ribbon of 70–200 serial sections. The serial sections were imaged by a field emission scanning electron microscope (JSM-IT800SHL; JEOL) with the Array Tomography Supporter software (System in Frontier). Imaging was done at 3 kV accelerating voltage, 268 pA beam current, 2,560 × 1.920 frame size, 6.5 mm working distance, 32.0 × 24.0 µm field of view (pixel size is 12.5 nm) and 2.67 µs dwell time, using the Scintillator Backscattered Electron Detector.

#### Quantification of MERCS size in electron micrographs

Electron micrographs were manually annotated using PHILOW software^55^. Mitochondria and ER in the vicinity of mitochondria were annotated and ER regions within 3 pixel (= 23.4 nm) of the mitochondrial periphery were defined as MERCS. One cell in the *Pdzd8* cKO condition was excluded from the statistical analysis because it was considered an outlier in the ROUT test (Q = 0.1%).

#### Quantification of MCI in serial electron micrographs of array tomography

The electron micrographs were aligned using the Linear Stack Alignment with Scale Invariant Feature Transform (SIFT) Plugin, implemented in ImageJ (NIH). Mitochondria, whose entire volume is within the imaging volume, were semi-automatically annotated using Empanada software^56^ and PHILOW software^55^. The volumes of mitochondria were calculated as the number of mitochondrial voxels multiplied by the voxel size (12.5 nm × 12.5 nm × 50 nm). The method for calculating mitochondrial surface area from electron microscopy images is as follows: The number of voxels along the circumference of mitochondrial cross-sections in the xy-plane was counted for each z-plane, and the totals were summed. This allowed for the enumeration of the voxels facing the surface in the direction perpendicular to the z-axis. Subsequently, this sum was multiplied by the 12.5 nm × 50 nm. To account for the mitochondrial surfaces facing along the z-axis the aforementioned voxel count was multiplied by 0.5, and the product was then multiplied by the dimensions of 12.5 nm × 12.5 nm. The sum of these two values was considered the total mitochondrial surface area. MCI was calculated by the following formula proposed by Vincent et al.^57^:

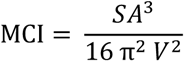

where SA is surface area and V is volume.

4, 9, 2, and 2 mitochondria in the control, *Pdzd8* cKO, *Fkbp8* KD, and *Pdzd8* cKO *+ Fkbp8* KD condition were excluded from the statistical analysis because it was considered outliers in the ROUT test (Q = 0.1%). 3D visualization of the mitochondrial structure was performed using Imaris version 9.6.0 (Bitplane).

#### Lentivirus production

Recombinant lentiviruses were produced as previously reported.^58^ 293T cells (BRC) were co-transfected with shuttle vectors (FUW-CreERT2-P2A-NeoR), HIV-1 packaging vectors Delta8.9 and VSV-G envelope glycoproteins, or shuttle vectors (pLKO-shFKBP8 or pLKO-scramble), LP1, LP2, and VSV-G using FuGENE transfection reagent (Promega, catalog no. E2311). Twenty-four hours after transfection, the media were exchanged with 8 mL of fresh DMEM supplemented with 10% FBS, and 1% penicillin-streptomycin, and 24 hours later, supernatants were harvested, spun at 500 × g to remove debris and filtered through a 0.45 μm filter (Sartorius). The filtered supernatant was concentrated to 125 μL using an Amicon Ultra-15 (molecular weight cut-off 100 kDa) centrifugal filter device (Merck Millipore), which was centrifuged at 4,000 × g for 60 minutes at 4°C. Then, 100 μL of viral supernatants was added to each 6-well dish containing MEFs.

#### Live cell imaging

Prior to imaging, cells were washed twice with PBS and then incubated in phenol red-free full DMEM supplemented with 10% FBS and 1% P/S for approximately 30 minutes. During the imaging, cells were maintained at 37°C in an incubation chamber (Tokai Hit). Images were acquired on a Nikon Ti2 Eclipse microscope with an A1R confocal, a CFI Plan Apochromat Lambda D 100X Oil (NA 1.45), a laser unit (Nikon; LU-N4, 405, 488, 561, and 640 nm), and filters (450/50nm, 525/50nm, 595/50nm for 405 nm, 488 nm, and 561 nm, respectively). All equipment was controlled via Nikon Elements software. Optical sectioning was performed at Nyquist for the longest wavelength. The resulting images were deconvoluted with NIS-elements (Nikon). The resulting images were processed with NIS-elements (Nikon) or ImageJ (NIH).

#### *HaloTag staining* (other than for single molecule tracking)

*PDZD8*-HaloTag knocked-in HeLa cells were incubated with 200 nM Janelia Fluor (JF) 549 dye (Promega, catalog no. GA1110) at 37°C o/n. 3 hours before the confocal imaging, 3 times PBS washes were performed and mediums were changed to JF549 free DMEM, then incubated at 37°C.

### Single particle tracking-photoactivation localization microscopy and analysis

#### Sample preparation

Coverslips (25 mm, No. 1.5, high tolerance, Warner Scientific) were cleaned according to a previously described protocol^59^ and stored in dry, sterile, 35mm tissue culture dishes sealed with parafilm until use within 3 months. Immediately before plating, coverslips were coated with 500 μM phenol red-free Matrigel (Corning) for one hour at 37°C. Coverslips were then washed once with sterile PBS before being overlayed with 2 mL of complete, phenol red-free DMEM. Simultaneously, 1 × 10^6^ COS7 cells per sample were trypsinized and resuspended directly into a transfection cocktail made of 750 ng PrSS-mEmerald-KDEL, 500 ng mTagRFP-T2-Mito-7, and 250 ng of msPDZD8-HaloTag-N1 mixed with Fugene HD (Promega) according to the manufacturer’s specifications. Cells were incubated in the suspension for 15 minutes at 37 degrees, and then the entire mixture was plated onto the coverslip and incubated for 18-22 hours until imaging. Any cells that were not imaged before 24 hours post transfection were discarded. Immediately prior to imaging, coverslips were loaded into a custom imaging chamber, labeled for 1 minute with 10 nM PA-JF646^60^ in OptiMEM (Gibco), and washed excessively (at least 5 times) with 10 mL of sterile PBS. Washing was performed while simultaneously aspirating from the chamber, taking extreme care to never expose the cells directly to the air. Cells were then washed once with 10 mL of complete, phenol red-free medium and allowed to recover in 1 mL of complete, phenol red-free DMEM for 15 minutes before imaging.

#### Microscopy and Imaging Conditions

Single molecule imaging was performed using a custom inverted Nikon TiE scope outfitted with a stage top incubation system to maintain cells at 37°C with 5% CO_2_ and appropriate humidity during imaging (Tokai Hit). Regions amenable to sptPALM (primarily the flat lamella of cells, 500 nm thick or less) were located by eye using the fluorescence of the ER label. The experimenter was always blinded to the single molecule tracers when choosing the cell region for imaging. Once a region with sufficiently flat ER was chosen, excitation was achieved using three fiber couples solid state laser lines (488 nm, 561 nm, 642 nm, Agilent Technologies) to illuminate the sample. The excitation beams were introduced into the system with a traditional rear-mount TIRF illuminator, which was used to manually set the angle of incidence beneath the critical angle to provide the most even illumination across the ER in the field of view. The illumination by the 488 nm and 561 nm lines were adjusted for each sample to minimize the bleed through into the single molecule channel, but both were always kept beneath 50 μW (488 nm) and 150 μW (561 nm) total power into the back aperture. The single molecule channel was always collevted with a constant 11.5 mW of 642 nm light introduced into the back aperature.

Emission light was collected using a 100x α-plan apochromat 1.49 NA oil immersion objective (Nikon Instruments). The collected light was focused onto three simultaneously running, electronically synchronized iXon3 electron multiplying charged coupled device cameras (EM-CCD, DU-897; Andor Technology), using a MultiCam optical splitter (Cairn Research) and sequential 565LP and 647LP dichroic mirrors (Chroma) within the optical path. The three emission paths were additionally cleaned up with a 525/50 BP, 605/70 BP, and 705/60 BP filter (Chroma) to filter extra light in the system.

The microscope was operated in sptPALM mode using only 128 × 128 pixel region on the camera (20.48 μm × 20.48 μm) to drive the system quickly enough to unambiguously track single proteins. The location of the region was always selected close to the center of the camera chip, since the objective being used is only chromatically well corrected at the center of the field of view. Imaging was performed with 5 msec exposure times, and the final speed was monitored using an oscilloscope directly coupled to the system (mean frame rate ∼95 Hz).

#### Trajectory generation and analysis

Single molecule localizations were linked to form trajectories using the TrackMate plugin in Fiji. Linking parameters were experimentally selected for each data set to minimize visible linkage artifacts an identified by eye. The resulting trajectories were then projected on to the ER network and manually curated for linkages that are close in 2D space but prohibitively far in the underlying ER structure itself. The resulting trajectories were exported from TrackMate and analyzed in Matlab for subsequent analysis, as described elsewhere.^30^

#### Correlative light and electron microscopy

Correlative light and electron microscopy was conducted as previously described.^61^ *PDZD8*-Halotag knocked-in HeLa cells were overexpressed with plasmids encoding Venus-FKBP8^N403K^ or YFP-ActA and then treated with 200 nM Janelia Fluor 549 dye (Promega) for 20 hours at 37°C. After that, cells were washed with PBS twice and plated in no. 1S gridded coverslip-bottom dishes (custom made, based on IWAKI 3922-035; coverslips were attached inversed side), precoated with carbon by a vacuum coater and then coated with poly-d-lysine (Merck, catalog no. P0899). The cells were fixed with 2% paraformaldehyde (Electron Microscopy Sciences) in PBS at room temperature for 10 minutes and then washed with PBS. Fluorescence imaging was conducted using Nikon AX confocal microscopy with a Nikon Spatial Array Confocal (NSPARC) detector and a CFI Plan Apochromat Lambda D 100X Oil (NA 1.45). The wavelength range of 503-545 nm or 582-618nm was obtained by excitation with a 488 nm or 561nm laser, respectively. The cells were then fixed with 2.5% glutaraldehyde in 0.1 M phosphate buffer (pH 7.4) for 2 days at 4°C. After washing with 0.1 M phosphate buffer, the cells were post-fixed with 1% OsO_4_ (Electron Microscopy Sciences), 1.5% potassium ferrocyanide (Fujifilm Wako pure chemical corporation, catalog no. 161-03742) in a 0.05 M phosphate buffer for 30 minutes. After being rinsed 3 times with H_2_O, the cells were stained with 1% thiocarbohydrazide (Sigma-Aldrich) for 5 minutes. After being rinsed with H_2_O for three times, the cells were stained with 1% OsO_4_ in H_2_O for 30 minutes. After being rinsed with H_2_O for two times at room temperature and three times with H_2_O at 50°C, the cells were treated with Walton’s lead aspartate (0.635% lead nitrate (Sigma-Aldrich), 0.4% aspartic acid (pH 5.2, Sigma-Aldrich)) at 50°C for 20 minutes. The cells dehydrated with an ascending series of ethanol (15 minutes each in 50% on ice, 70% on ice, and 10 minutes each in 90%, 95% ethanol/H_2_O at room temperature, 10 minutes in 100% ethanol 4 times at room temperature) were embedded in epoxy resin (LX-112) by covering the gridded glass with a resin-filled beam capsule. LX-112 resin was made by mixing 4.85 g of NMA (Ladd Research Industries), 7.8 g of LX-112 (Ladd Research Industries), 2.35 g of DDSA (Ladd Research Industries), and 0.3 g of BDMA (Ladd Research Industries). Polymerization was carried out at 42°C for 12 hours and 60°C for 72 hours. After polymerization, the gridded coverslip was removed and the resin block was trimmed to a square of about 150 to 250 μm. The block was sectioned using an ultramicrotome (EM UC7, Leica) equipped with a diamond knife (Ultra JUMBO 45 degree, DiATOME) to cut 50 nm thick sections. The serial ultra-thin sections were collected on the cleaned silicon wafer strip and imaged with a scanning electron microscope (JSM-IT800SHL; JEOL). Imaging of the Venus-FKBP8^N403K^ overexpressing cell was done at 2 kV accelerating voltage, 34.1 pA beam current, 5,120 × 3,840 frame size, 6.5 mm working distance, 12.8× 9.6 µm field of view (pixel size is 2.5 nm) and 1.33 µs dwell time, using the Scintillator Backscattered Electron Detector. Imaging of the YFP-ActA-overexpressing cell was done at 2 kV accelerating voltage, 48.8 pA beam current, 2,560 × 1,920 frame size, 6.9 mm working distance, 12.8× 9.6 µm field of view (pixel size is 5 nm) and 14.1 µs dwell time, using the Scintillator Backscattered Electron Detector. The images taken by confocal microscopy were processed with ImageJ (NIH). The electron micrographs were stitched by Stitch Sequence of Grids of Images Plugin and aligned using the Linear Stack Alignment with Scale Invariant Feature Transform (SIFT) Plugin, implemented in ImageJ (NIH). After that, pixel size of images from the Venus-FKBP8-overexpressing cell was converted 2.5 nm to 5 nm by OpenCV’s resize function. Mitochondria and ERs in the vicinity of mitochondria in electron micrographs were semi-automatically annotated using Empanada software^56^ and PHILOW software^55^. ER regions within 5 pixels (= 25 nm) of the mitochondrial periphery were defined as MERCS. Reconstructing segmented images of the electron micrographs to 3-dementional images and overlaying it with fluorescence images were conducted using Imaris software (Bitplane). The z projection of electron micrographs was created using ImageJ (NIH). For the analysis of mitochondrial membrane curvature at PDZD8-localized MERCS, the arbitrary area was extracted from the z-projection images of 3D-reconstructed electron micrographs merged with fluorescence images and counted the number of PDZD8-localized MERCS with positive, negative, and neutral OMM using ImageJ.

#### Cryo-Correlative light and electron microscopy

Cells were transiently transfected with FKBP8^N403K^-mScarlet plasmid for 24 hours and then were transferred to cryo-EM grids (Quantifoil, 200-Au-mesh with carbon film) (pre-treated with 50 ug/ml fibronectin for at least 15 minutes) and incubated for 3 hours. The cells were incubated with 5% glycerol in the media for 15 minutes right before freezing after which the media was exchanged with PBS and 5% glycerol, and the grids were plunge-frozen. A Leica GP2 was used for plunge freezing with double blotting each for 7 seconds. The grids were stored in liquid nitrogen for the following steps. First, the grids were imaged using a Zeiss LSM900 AiryScan and the Linkam cryo-stage to screen for overall quality and transfection efficiency. Second, 4 grids were milled in one session using an Aquilos2 FIB-SEM equipped with the Delmic IceShield. 35 lamellae were generated with a thickness of around 170 nm and varying surface areas. These lamellae were loaded on a Titan Krios equipped with a K3 detector and BioQuantum energy filter. High-resolution medium-mag montages of lamellae were collected and manually inspected for MERCS. MERCS were then targeted for high-resolution tilt series acquisition at a pixel size of 2.07 Å/pixel. Grids were loaded into the cassette with a lamella pre-tilt of −9 degrees, thus the tilt series acquisition started at 9 degrees, with a target range of −42 to +60 degrees on the grid (−51 to +51 on the lamellae) acquired in 3-degree increments in a dose-symmetric fashion using SerialEM.^62^ The data collection was monitored live using Warp.^63^ After cryo-ET data collection, the lamellae were imaged in the Zeiss AiryScan with a 100x objective. This was feasible since we discovered that the mScarlet tag survives TEM radiation. The fluorescence images were overlaid on the TEM montages using the MAPS software (Thermo Fisher Scientific). Tilt series alignment was done using AreTomo. Weighted back projection tomogram reconstructions were CTF-deconvolved using ISONET.^64^ Initial Segmentation was done using TomoSegMemTV^65^ on the deconvoluted tomograms. The segmentations were corrected and labelled manually using Amira. The segmentations were subsequently processed and analyzed using the surface morphometrics analysis toolkit.^36^ In the OE condition, 73 out of 116 collected tomograms (63%) contained at least one contact site, while in the control, only 19 out of 49 collected tomograms (38%) contained at least one contact site. Full analysis was on 20 MERCS for the OE condition, and 6 for the control.

#### Sample preparation and LC-MS/MS for Pdzd8-3×HA knock-in mouse brain

Pdzd8-3×HA knock-in mice and control littermates at postnatal 10 days were put to sleep using medetomidine hydrochloride (Domitor, Nippon zenyaku kogyo, 0.75 mg/kg), midazolam (Sandoz, 4 mg/kg) and butorphanol (Vetorphale, Meiji Seika Pharma Co., Ltd, 5 mg/kg). Pups were then put on the ice for 5 minutes and exsanguinated by terminal intracardial perfusion with ice-cold 2% paraformaldehyde (Merck) in phosphate-buffered saline (PBS). The neocortex was then removed and sonicated five times for 30 seconds with ice-cold lysis buffer (20 mM Hepes-NaOH pH7.5, 1 mM EGTA, 1 mM MgCl2, 150 mM NaCl, 0.25% Na-deoxycholate, 0.05% SDS, 1% NP40, Benzonase (Merck), PhosSTOP phosphatase inhibitor (Roche) and cOmplete protease inhibitor cocktail (Roche)). After the lysates were centrifuged at 20,000 × g for 15 min at 4°C, the resulting supernatants were incubated for 3 hours at 4°C with a 2.5 µL slurry of Sera-Mag SpeedBeads Protein A/G (Cytiva) pre-incubated with 2.5 µL of anti-HA-tag rabbit mAb (Cell signaling technology, C29F4). The beads were washed four times with the lysis buffer and then twice with 50 mM ammonium bicarbonate. Proteins on the beads were digested by adding 200 ng trypsin/Lys-C mix (Promega) at 37 °C overnight. The resulting digests were reduced, alkylated, acidified, and desalted using GL-Tip SDB (GL Sciences). The eluates were evaporated and dissolved in 0.1% trifluoroactic acid and 3% ACN. LC-MS/MS analysis of the resultant peptides was performed on an EASY-nLC 1200 UHPLC connected to an Orbitrap Fusion mass spectrometer through a nanoelectrospray ion source (Thermo Fisher Scientific). The peptides were separated on a C18 reversed-phase column (75 mm [inner diameter] x 150 mm; Nikkyo Technos) with a linear 4%–32% ACN gradient for 0–100 minutes, followed by an increase to 80% ACN for 10 minutes and final hold at 80% ACN for 10 minutes. The mass spectrometer was operated in data-dependent acquisition mode with a maximum duty cycle of 3 seconds. MS1 spectra were measured with a resolution of 120,000, an automatic gain control (AGC) target of 4e5, and a mass range of 375–1,500 m/z. HCD MS/MS spectra were acquired in the linear ion trap with an AGC target of 1e4, an isolation window of 1.6 m/z, a maximum injection time of 35 ms, and a normalized collision energy of 30. Dynamic exclusion was set to 20 s. Raw data were directly analyzed against the SwissProt database restricted to Mus musculus using Proteome Discoverer version 2.5 (Thermo Fisher Scientific) with Sequest HT search engine for identification and label-free precursor ion quantification. The search parameters were as follows: (i) trypsin as an enzyme with up to two missed cleavages; (ii) precursor mass tolerance of 10 ppm; (iii) fragment mass tolerance of 0.6 Da; (iv) carbamidomethylation of cysteine as a fixed modification; and (v) acetylation of the protein N-terminus and oxidation of methionine as variable modifications. Peptides and proteins were filtered at a false discovery rate (FDR) of 1% using the Percolator node and Protein FDR Validator node, respectively. Label-free quantification was performed on the basis of the intensities of precursor ions using the Precursor Ions Quantifier node. Normalization was performed such that the total sum of abundance values for each sample over all peptides was the same.

#### Sample preparation and LC-MS/MS for PDZD8-TurboID KI HeLa cells

*PDZD8*-TurboID knocked-in HeLa cells were plated into a 15 cm dish at the density of 2×10^6^ cells/dish and cultured two overnight. The cells were treated with Biotin at the final concentration of 50 μM, and incubated for 6 hours. The cells were washed twice with ice-cold Hepes-saline and lysed in 6 M guanidine-HCl (Wako) containing 100 mM HEPES-NaOH (pH7.5), 10 mM TCEP (Nacalai), and 40 mM chloroacetamide (Sigma). Following sample preparation and analysis were conducted as described previously.^66^ Briefly, after heating and sonication, proteins (1.3 mg each) were purified by methanol–chloroform precipitation and resuspended in 200 μL of PTS buffer (12 mM SDC, 12 mM SLS, 100 mM Tris-HCl, pH8.0). After sonication and heating at 95 °C for 10 minutes, the protein solutions were diluted 5-fold with 100 mM Tris-HCl, pH8.0 and digested with 13 μg of trypsin (Pierce) at 37°C overnight. After heating at 95 °C for 10 minutes, the digested peptides were incubated with the ACN-prewashed Tamavidin 2-REV beads (FUJIFILM Wako) for 3 hours at 4°C. After washing five times with TBS (50 mM Tris-HCl, pH7.5, 150 mM NaCl), biotinylated peptides were eluted for 15 minutes at 37°C twice with 100 µL of 1 mM biotin in TBS. The combined eluates were desalted using GL-Tip SDB, evaporated, and redissolved in 0.1% TFA and 3% ACN. LC-MS/MS analysis of the resultant peptides was performed on an EASY-nLC 1200 UHPLC connected to an Orbitrap Fusion mass spectrometer. The peptides were separated on the C18 reversed-phase column with a linear 4%–32% ACN gradient for 0–60 minutes, followed by an increase to 80% ACN for 10 minutes and final hold at 80% ACN for 10 minutes. The mass spectrometer was operated in data-dependent acquisition mode with a maximum duty cycle of 3 seconds. MS1 spectra were measured with a resolution of 120,000, an AGC target of 4e5, and a mass range of 375–1,500 m/z. HCD MS/MS spectra were acquired in the linear ion trap with an AGC target of 1e4, an isolation window of 1.6 m/z, a maximum injection time of 200 ms, and a normalized collision energy of 30. Dynamic exclusion was set to 10 s. Raw data were directly analyzed against the SwissProt database restricted to Mus musculus using Proteome Discoverer version 2.5 with Sequest HT search engine for identification and label-free precursor ion quantification. The search parameters were as follows: (i) trypsin as an enzyme with up to two missed cleavages; (ii) precursor mass tolerance of 10 ppm; (iii) fragment mass tolerance of 0.6 Da; (iv) carbamidomethylation of cysteine as a fixed modification; and (v) acetylation of the protein N-terminus, oxidation of methionine, and biotinylation of lysine as variable modifications. Peptides and proteins were filtered at a FDR of 1% using the Percolator node and Protein FDR Validator node, respectively. Label-free quantification was performed on the basis of the intensities of precursor ions using the Precursor Ions Quantifier node. Normalization was performed such that the total sum of abundance values for each sample over all peptides was the same.

### Sample preparation and LC-MS/MS for Proximity labeling-Mass spectrometry for PDZD8-TurboID KI Neuro2a cells

#### in situ Biotinylation

*Pdzd8*-v5-TurboID knock-in cell (mouse Neuro2a) were grown in T75 (75cm^2^) cell culture flask. One flask per each triplicate samples were prepared for MS analysis. The cells were then treated with biotin (50 μM) in complete DMEM medium for 30 minutes at 37°C. The labeled cells were washed three times with cold DPBS 30 minutes after the in situ biotinylation, following next cell lysis. The cells were lysed with 750 μl of 1 × TBS (25 mM Tris, 0.15 M NaCl, pH 7.2) containing 2% SDS and 1 × protease inhibitor cocktail. Lysates were clarified by ultrasonication (Bioruptor, diagenode) to physically break down nucleic acid such as DNA for 5 minutes three times with iced water bath.

#### Enrichment of biotinylated peptide and LC-MS/MS

The 4 ml of cold acetone were added to the lysates and incubated at −20°C for at least 2 hours to 16 hours. After first precipitation with acetone, the samples were centrifuged at at 13,000 × *g* for 10 minutes at 4°C and the supernatant was gently discarded. To completely reconstitute all proteins, samples were incubated with cooled 10% TBS/90% acetone at −20°C for at least 2 hours to 16 hours. The samples were centrifuged at at 13,000 ×*g* for 10 minutes at 4°C and the supernatant was gently discarded. The pellets were reconstituted with 500 μl of 8 M urea containing 50 mM ABC, followed by measuring proteins concentration using BCA assay. Denaturation was performed at 650 rpm for 1 hour at 37°C. Samples were reduced by reducing agent, 10 mM DTT and alkylated with 40 mM IAA at 650 rpm for 1 hour at 37°C, respectively. The. Samples were diluted 8-fold with 50 mM ABC containing 1 mM CaCl_2_. Samples were digested with (50:1 w/w) trypsin at 650 rpm for at least 6 to 18 hours at 37°C. Insoluble fractions were removed by centrifugation for 10 minutes at 10,000 × *g*. The 150 μl of Streptavidin magnetic beads were firstly washed with 1× TBS containing 2 M urea four times and then added to the digested solution. The binding with the beads were for 1 hour room temperature, followed by washing the beads two times with 50 mM ABC containing 2 M urea. The washed beads were washed with pure water compatible with LC-MS and transferred to new protein lobind tubes (Eppendorf). For elution of biotinylated peptides from the beads, 150 μl of elution solution (80% ACN, 20% pure water, 0.2% TFA, and 0.1% formic acid) were incubated at 60°C three times repeatedly. Total elution fractions were completely dried using a speedvac. The resulting peptides were analyzed by Q Exactive Plus orbitrap mass spectrometer (Thermo Fisher Scientific, MA, USA) equipped with a nanoelectrospray ion source. To separate the peptide mixture, we used a C18 reverse-phase HPLC column (500 mm × 75 μm ID) using an acetonitrile/0.1% formic acid gradient from 4 to 32.5% for 120 minutes at a flow rate of 300 nL/min. For MS/MS analysis, the precursor ion scan MS spectra (m/z 400∼2000) were acquired in the Orbitrap at a resolution of 70,000 at m/z 400 with an internal lock mass. The 15 most intensive ions were isolated and fragmented by High-energy collision induced dissociation (HCD).

#### LC-MS/MS data processing of biotinylated peptides

All MS/MS samples were analyzed using the Sequest Sorcerer platform (Sagen-N Research, San Jose, CA, USA Sequest was set up to search the Mus musculus protein sequence database (86320 entries, UniProt, http://www.uniprot.org/) assuming the digestion enzyme stricttrypsin. Sequest was searched with a fragment ion mass tolerance of 1.00 Da and a parent ion tolerance of 10.0 ppm. Carbamidomethylation of cysteine was specified in Sequest as a fixed modification. Oxidation of methionine, acetyl of the n-terminus, phospho of serine, threonine and tyrosine and biotin of lysine were specified in Sequest as variable modifications. Scaffold (Version 4.11.0, Proteome Software Inc., Portland, OR, USA) was used to validate MS/MS-based peptide and protein identifications. Peptide identifications were accepted if they could be established at greater than 95.0% probability by the Scaffold Local FDR algorithm. Protein identifications were accepted if they could be established at greater than 99.0% probability and contained at least 2 identified peptide. Protein probabilities were assigned by the Protein Prophet algorithm.^67^ Proteins that contained similar peptides and could not be differentiated based on MS/MS analysis alone were grouped to satisfy the principles of parsimony. Proteins were annotated with Gene Ontology (GO) terms from the National Center of Biotechnology Information database (NCBI; downloaded November 1, 2019).^68^

---

**Extended Data Fig 1.**
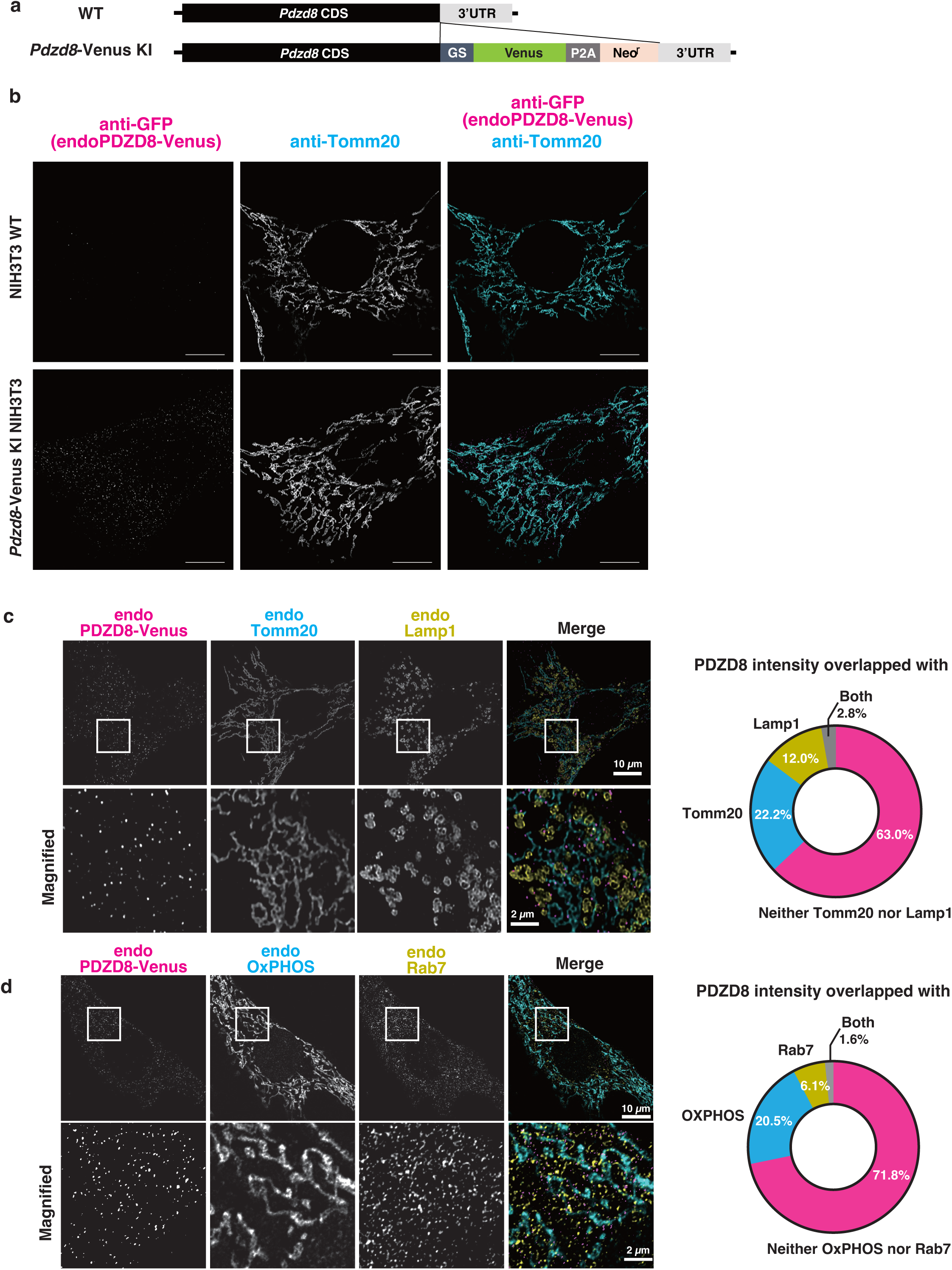
PDZD8 localizes in the close proximity of mitochondria, lysosome, and late endosome. **a,** Diagram describing the genomic locus of *Pdzd8* in the *Pdzd8*-Venus KI NIH3T3 cell. The sequence for Venus-P2A-Neomycin resistance gene (Neo^r^) was knocked-in at the C-terminus of the *Pdzd8* coding sequence. **b,** Validation of endogenous PDZD8-Venus immunostaining of *Pdzd8*-Venus KI NIH3T3 cells. The cells were stained with antibodies to GFP, to Tomm20, and to Lamp1 (not shown). Note that the signal of anti-GFP was observed in the KI cell, but not in the WT cell. Scale bars, 10 μm. **c, d,** Analyses of PDZD8 localization by immunofluorescence using *Pdzd8*-Venus KI NIH3T3 cells. The cells were stained with antibodies to GFP, to Tomm20, and to Lamp1 (**c**), or with antibodies to GFP, to OXPHOS complex, and to Rab7 (**d**). The boxed regions of the top panels are shown at higher magnification in the corresponding lower panels. Scale bars, 10 μm or 2 μm as indicated. Data are representative of two independent experiments. Pie charts show the percentage of endogenous PDZD8-Venus intensity overlapping with areas positive for both Tomm20 and Lamp1, Tomm20-only, Lamp1-only, or the other areas of images (**c**), or with areas positive for both OXPHOS and Rab7, OXPHOS -only, Rab7-only, and the other areas of images (**d**).

**Extended Data Fig 2.**
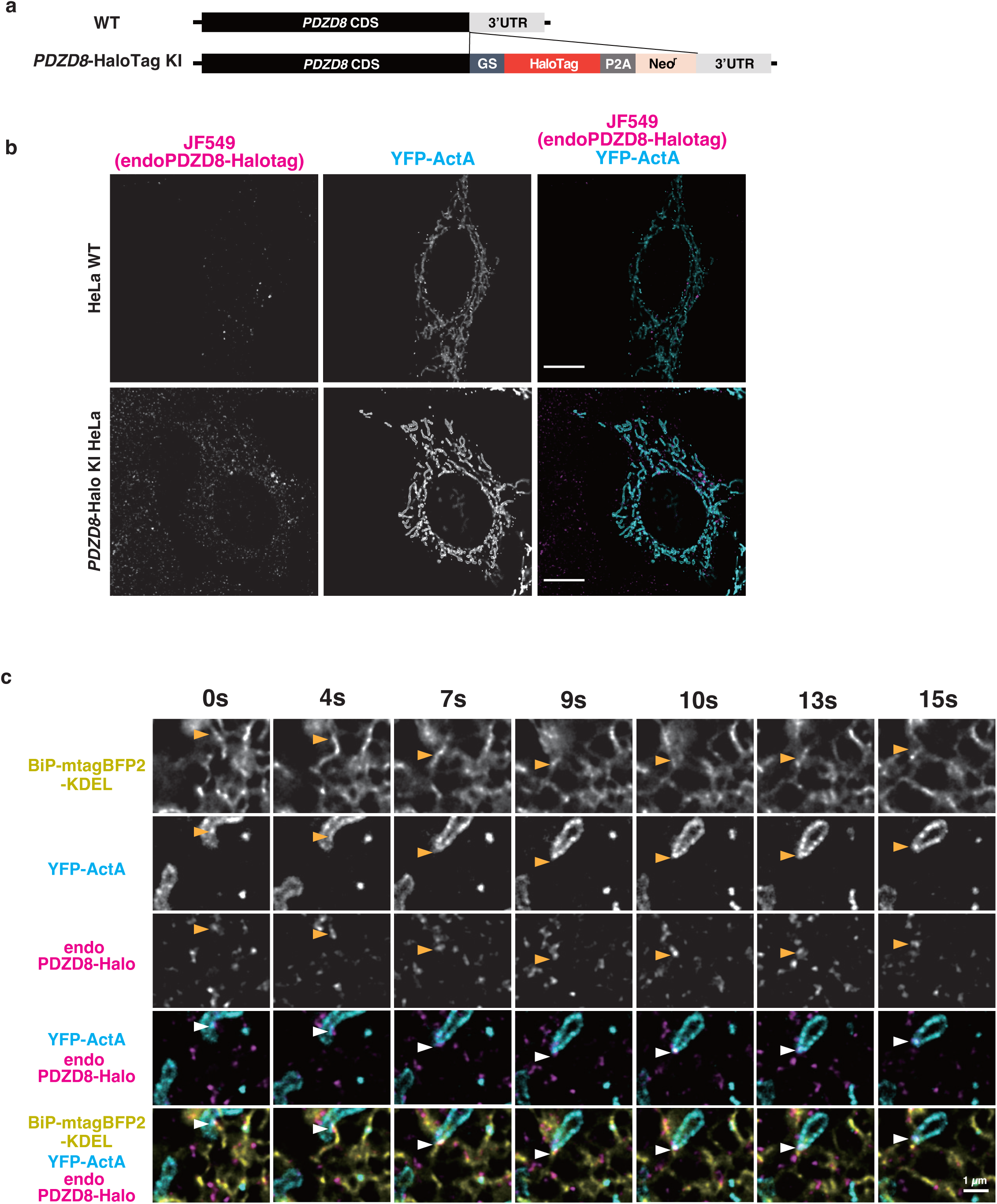
Endogenous PDZD8 localizes in the close proximity of mitochondria in living cells. **a,** Diagram describing the genomic locus of PDZD8 in the *PDZD8*-Halotag KI HeLa cell. The sequence for Halotag-P2A-Neo^r^ was knocked-in at the C-terminus of the *PDZD8* coding sequence. **b,** Snap shots from live imaging of wild-type HeLa cells and *PDZD8*-HaloTag KI HeLa cells transiently transfected with YFP-ActA (cyan). Endogenous PDZD8-HaloTag was labeled with Janelia Fluor 549 dye (magenta). Scale bars, 10 μm. **c,** Representative time-lapse images from a live-cell imaging of a *PDZD8*-HaloTag KI HeLa cell transiently transfected with an ER-marker BiP-mtagBFP2-KDEL (yellow) and an OMM-marker YFP-ActA (cyan), and treated with Janelia Fluor 549 dye (magenta) for labeling endogenous PDZD8-HaloTag. The arrowheads indicate the puncta of endo-PDZD8-HaloTag colocalizing with mitochondria and ER. Note that the PDZD8 dot moved together with the ER-mitochondria contact site.

**Extended Data Fig 3.**
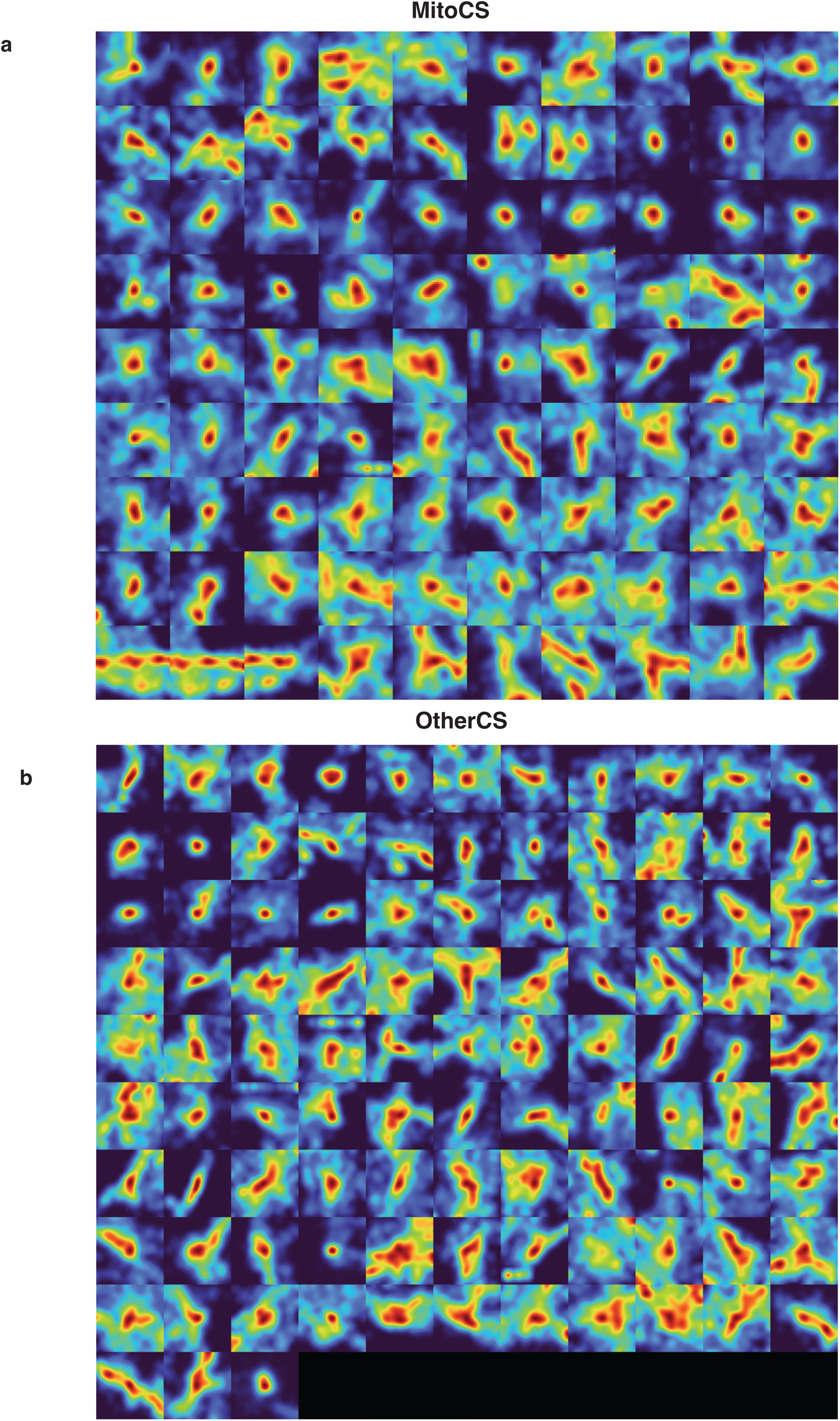
Spatially defined likelihood of PDZD8 localization in MitoCS or OtherCS. **a, b,** All MitoCS (**a**) and OtherCS (**b**) identified within the dataset are shown.

**Extended Data Fig 4.**
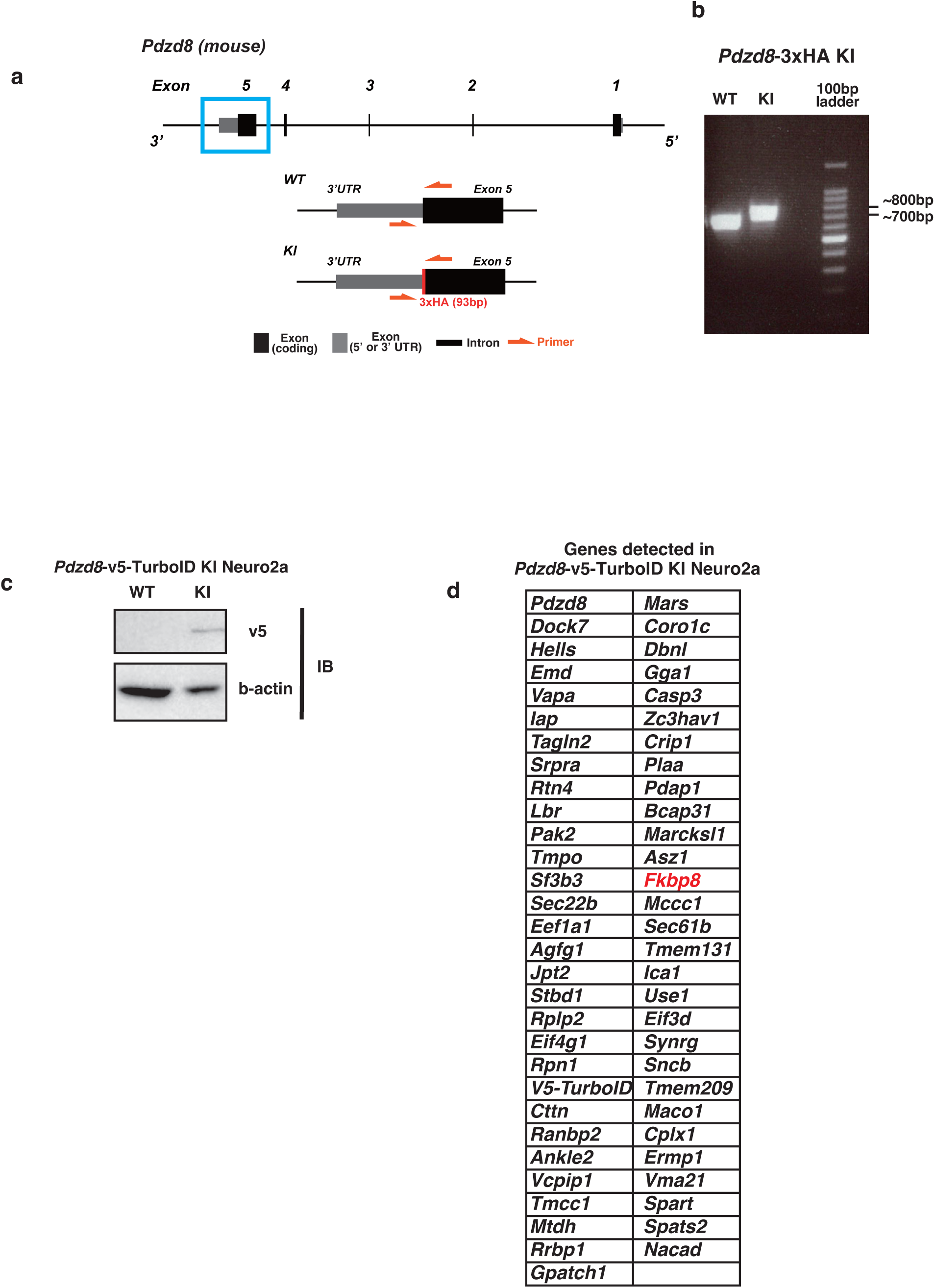
Establishment of *Pdzd8*-3×HA knock-in mice, the proteome of *Pdzd8*-v5-TurboID KI Neuro2a. **a,** Diagram showing genomic sequence of *Pdzd8*-3×HA knock-in mouse. The HA tag sequence was inserted in the end of *Pdzd8* coding region. **b,** Genomic analysis of *Pdzd8*-3×HA knock-in mouse. The sequence around the *Pdzd8* stop codon was amplified from genomic DNA of wild type (WT) or *Pdzd8*-3×HA knock-in (KI) mouse with primers as shown in (A). Primer sequence; 5’-gag gct tgc cga cag aag ag - 3’ and 5’-agt gag aca tca cac ata cac aaa −3’. **c,** Immunoblot analysis of WT and *Pdzd8*-v5-TurboID KI Neuro2a cells with antibodies to v5 and β-actin. **d,** Proteins detected by proximity labeling of *Pdzd8*-v5-TurboID KI Neuro2a cells incubated in 50 µM biotin-containing medium for both 3 hours and 16 hours. Note that FKBP8 is in this list.

**Extended Data Fig 5.**
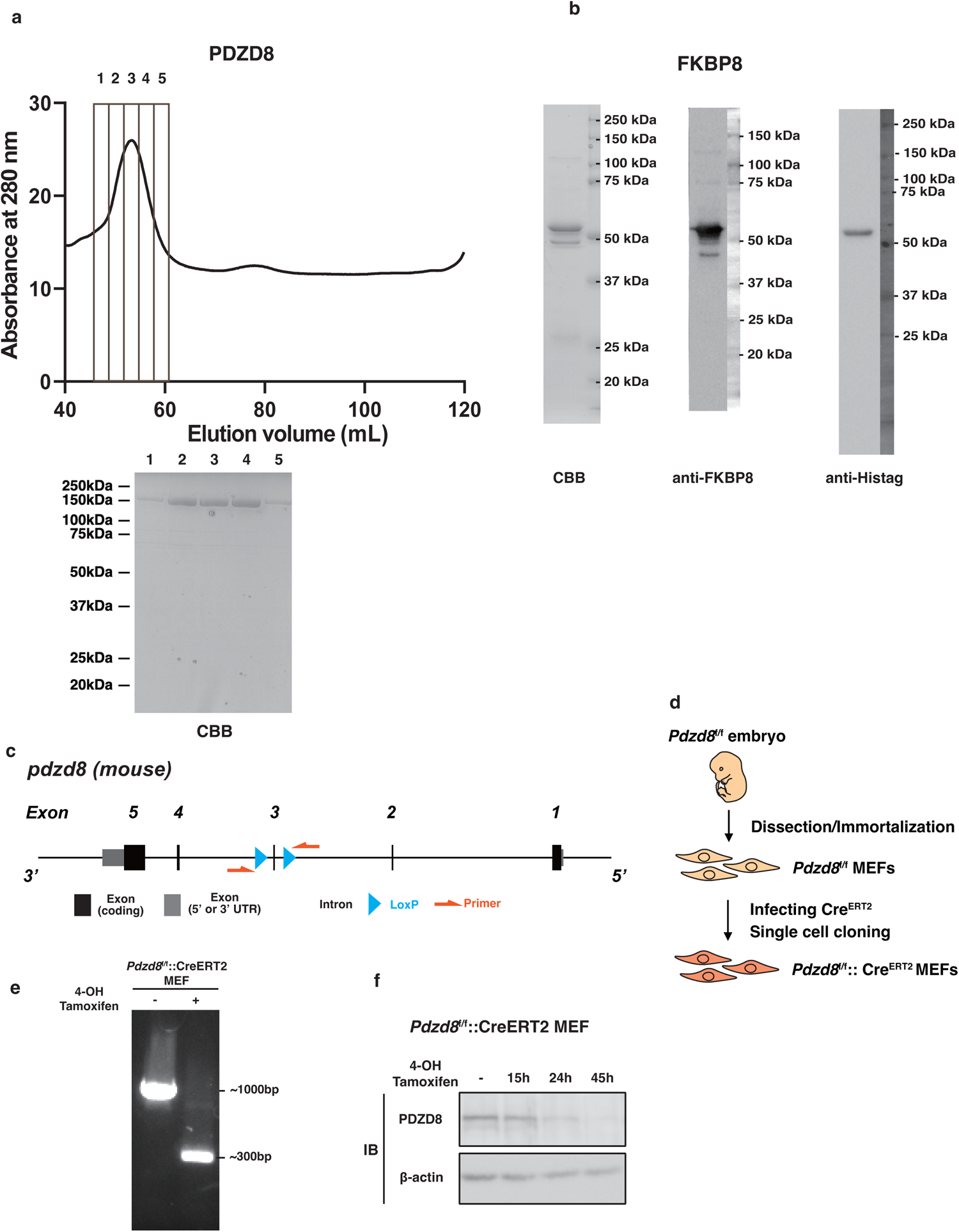
Properties of recombinant protein used for SPR and the establishment of *Pdzd8* ^f/f^::Cre^ERT2^ MEF. **a,** Purification profile of hPDZD8(1, 28-)-FLAG by size exclusion chromatography after anti-FLAG affinity column purification. Fractions 1 to 5 are used for SPR analyses. **b,** SDS-PAGE gel analysis and immunoblot analysis of the hFKBP8 (1-380) - Histag. Coomassie Brilliant Blue staining confirmed the purity of recombinant FKBP8 and immunoblot analysis with antibodies to Histag and FKBP8 showed the major band (upper one) is anti-HisTag-positive and major band (upper one) and minor band (lower one) are anti-FKBP8-positive, which suggests major band (upper one) is Full-length of FKBP8 (1-380) - Histag and, minor band (lower one) is FKBP8 whose C-terminal sequence is cleaved. **c,** Diagram showing genomic locus of *Pdzd8* conditional KO mouse*. Pdzd8* exon 3 is flanked by loxP sites. Cre-mediated recombination excises the flanked DNA, creating a frame-shift mutation which results in total knockout of the *Pdzd8* gene. **d,** Schematic diagram for generating tamoxifen-inducible *Pdzd8* conditional knock-out cell lines (*Pdzd8*^f/f^::Cre^ERT2^ MEFs). **e,** Genomic analysis of *Pdzd8*^f/f^::Cre^ERT2^ MEFs treated with mock or 1 μM of 4-hydroxy tamoxifen/ethanol for 24 hours. The sequence around exon 3 was amplified from genomic DNA of each cell lysate with primers as shown in **c**. Primer sequence; 5’-GCC AGT CAG AGA CCA TGA GAA A −3’ and 5’-ACA TCT GTT TTG TTT ACC ACT CTG C −3’. **f,** Immunoblot analysis of *Pdzd8*^f/f^::Cre^ERT2^ MEFs with antibodies to PDZD8 and β-actin. Cells were treated with mock (-) or 1 μM of 4-hydroxy tamoxifen for indicated time.

**Extended Data Fig 6.**
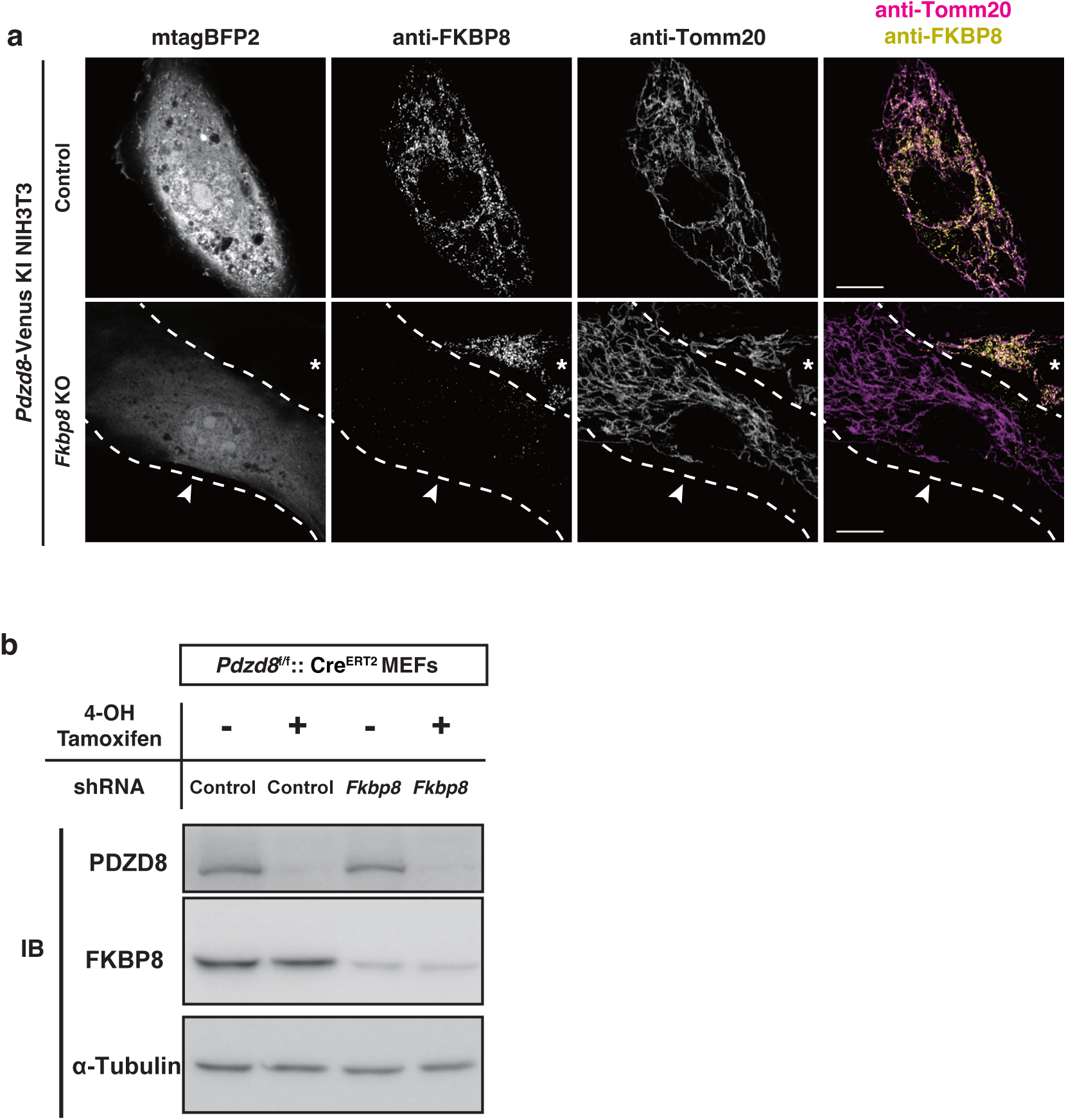
Validation of knocking out of FKBP8, knocking down of FKBP8, and conditional knocking out of PDZD8. **a,** Validation of knocking out of endogenous FKBP8. *Pdzd8*-Venus NIH3T3 cells transfected with the control gRNA (upper) or three gRNAs against FKBP8 (bottom) and transfection marker mtagBFP2, and stained with antibodies to FKBP8, and Tomm20 for visualizing endogenous FKBP8 (yellow) and mitochondrial outer membrane (magenta), respectively. Note that the signal of anti-FKBP8 was observed in the control cells, but not in *Fkbp8* KO cells. Arrowheads indicate the nucleus of the cell transfected. Asterisks indicate the nucleus of the cell not transfected. Scale bars, 5 μm. **b,** Immunoblot of *Pdzd8*^f/f^::Cre^ERT2^ MEFs infected with lentivirus carrying shControl or shFKBP8, treated with or without 0.5 µM 4-OHT. Cell lysates were subjected to immunoblotting with antibodies to PDZD8, FKBP8, and α-tubulin (bottom).

**Extended Data Fig 7.**
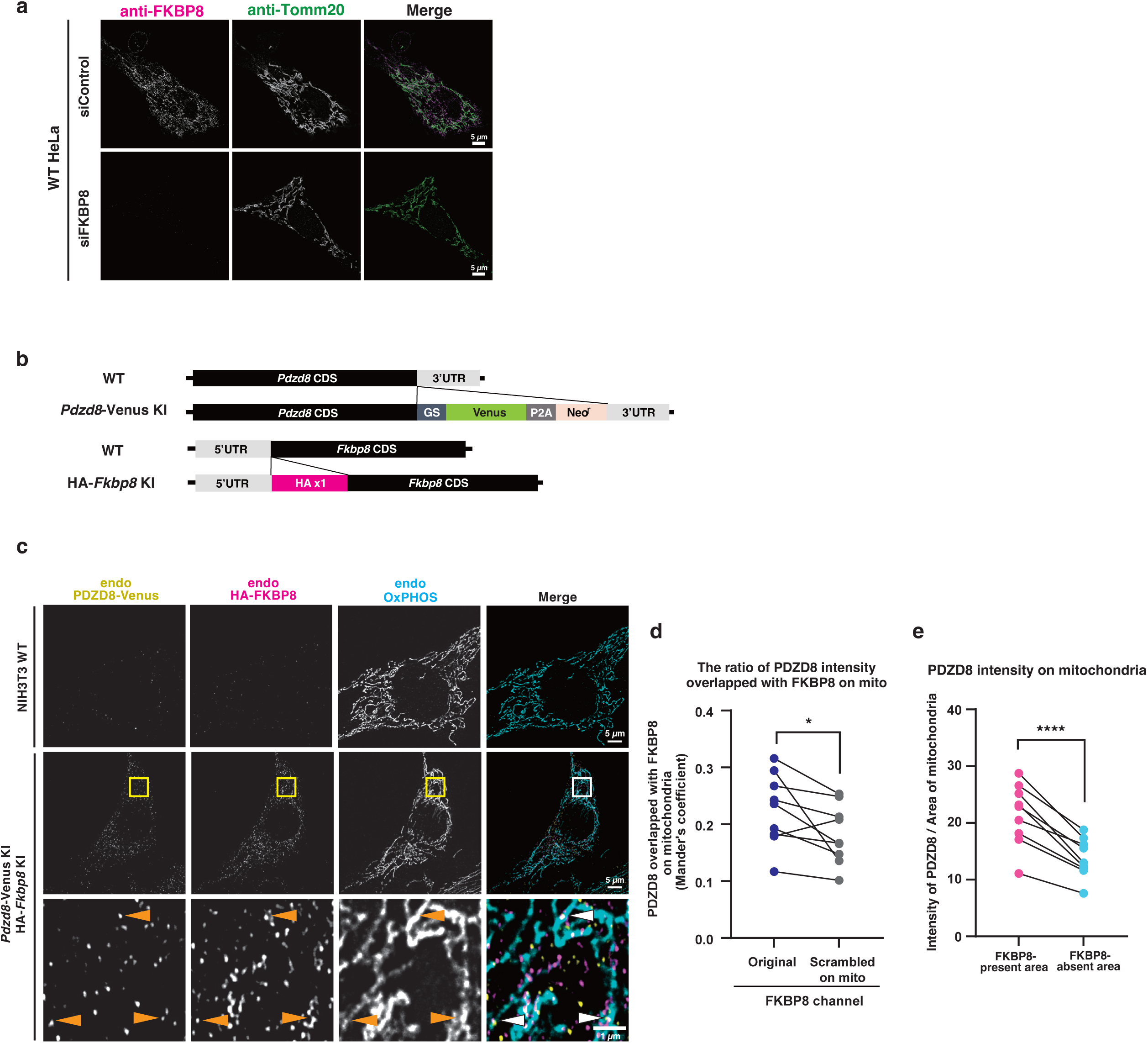
Validation of specificity of anti-FKBP8 antibody and colocalization analysis of PDZD8 and FKBP8 in NIH3T3 cells. **a,** Validation of specificity of the anti-FKBP8 antibody used in Figure 5a. HeLa cells transfected with siControl or siFKBP8 were stained with antibodies to FKBP8 and to Tomm20. Note that the signal of anti-FKBP8 was observed in siControl cells, but not in siFKBP8 cells. Scale bars, 5 μm. **b,** Diagram describing the genomic sequence of *Pdzd8*-Venus KI and HA-*Fkbp8* double KI (DKI) cells. The sequence of HA tag was knocked-in at the N-terminus of *Fkbp8* coding region in *Pdzd8*-Venus KI NIH3T3 cells. **c,** Immunofluorescence analysis of *Pdzd8*-Venus and HA-*Fkbp8* DKI cells. The cells were stained with antibodies to GFP, to HA, and to OXPHOS complex. The boxed regions of the top panels are shown at higher magnification in the corresponding middle panels. Arrowheads indicate PDZD8 colocalized with FKBP8 on mitochondria. Scale bars, 5 μm or 1 μm (higher magnification images). Data are representative of three independent experiments. **d,** The ratios of PDZD8 intensity overlapped with FKBP8 on mitochondria (Mander’s coefficients) were determined for images as described in **c**. The scrambled FKBP8 images were created by shuffling pixels within mitochondria in the FKBP8 channel. Data are representative of three independent experiments (9 cells). Paired t-test was used to test statistical significance. \**P* < 0.05. **e,** The means of PDZD8 intensity in the FKBP8-present or FKBP8-absent area on mitochondria were determined for images as in **c**. Data are representative of three independent experiments (9 cells). Paired t test was used to test statistical significance. \*\*\*\**P* < 0.0001.

**Extended Data Fig 8.**
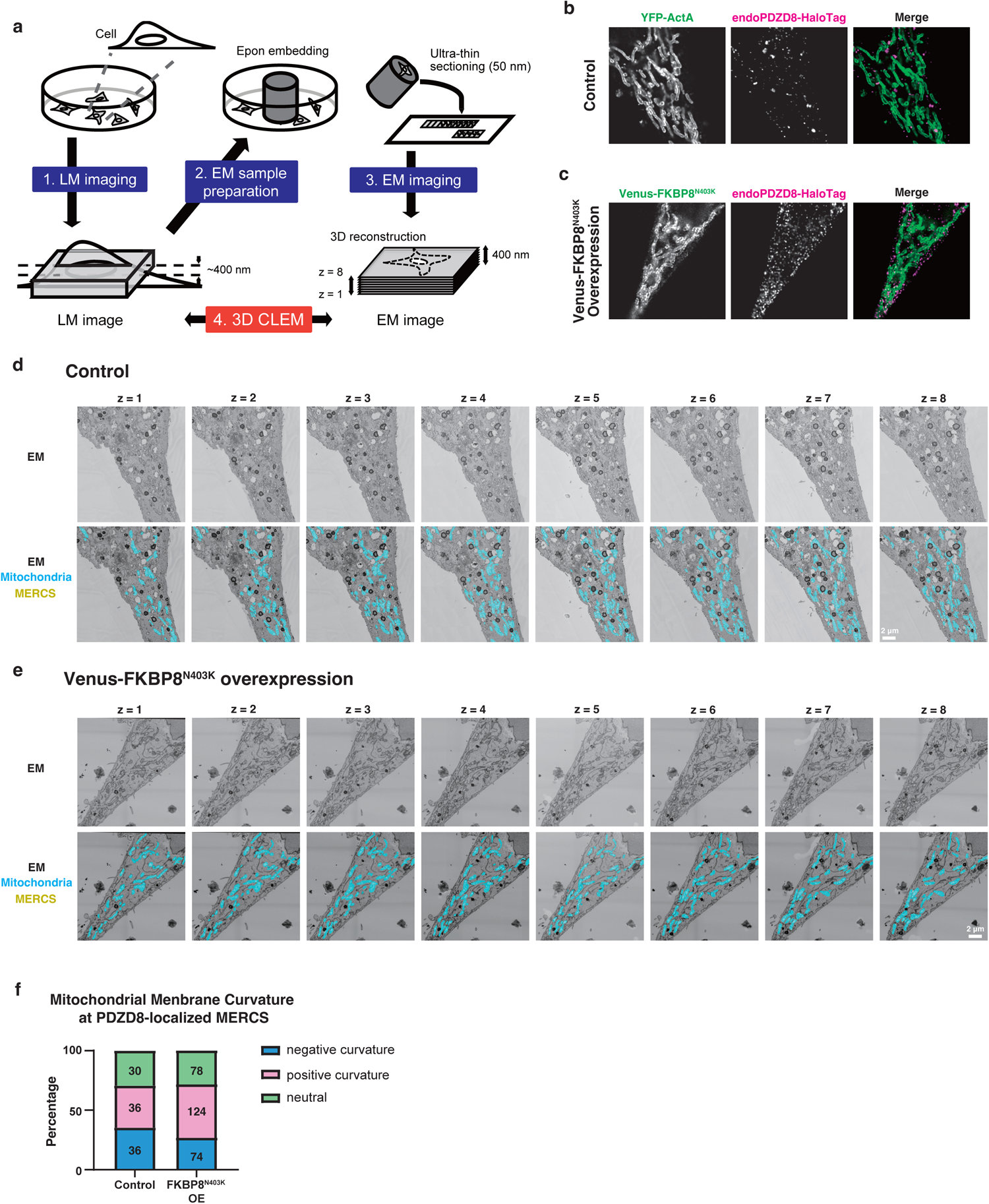
Electron micrographs of Correlative Light and Electron Microscopy (CLEM) analysis. **a,** Scheme of Correlative Light and Electron Microscopy (CLEM) analysis. *PDZD8*-HaloTag KI HeLa cells, which were labeled with JF549-conjugating HaloTag ligands and expressed with Venus-FKBP8^N403K^ or YFP-ActA as a control, were observed in confocal microscopy with a Nikon Spatial Array Confocal (NSPARC) detector after fixation. Subsequently, the same cells were imaged by field emission scanning electron microscope (FE-SEM). The resulting electron micrographs were utilized to reconstruct 3-dimensional (3D) images, and then the corresponding areas in the fluorescence image were re-identified in the electron micrographs. **b, c,** Fluorescence images of cells overexpressing Venus-FKBP8^N403K^ or YFP-ActA obtained by confocal microscopy with NSPARC detector. **d, e,** Electron micrographs of the serial 8 slices corresponding to the optical section of fluorescence images (indicated in **b** and **c**) of the control (**d**) and Venus-FKBP8^N403K^ OE cell (**e**). Mitochondria and the ER within 25 nm of mitochondria (MERCS) were indicated as cyan and yellow respectively. **f,** Quantification of mitochondrial membrane curvature at PDZD8-localized MERCS. The arbitrary area was extracted from the 3D-reconstructed images and counted the number of PDZD8-localized MERCS with positive, negative, and neutral OMM. The number of each type of MERCS was indicated in the graph.

**Extended Data Fig 9.**
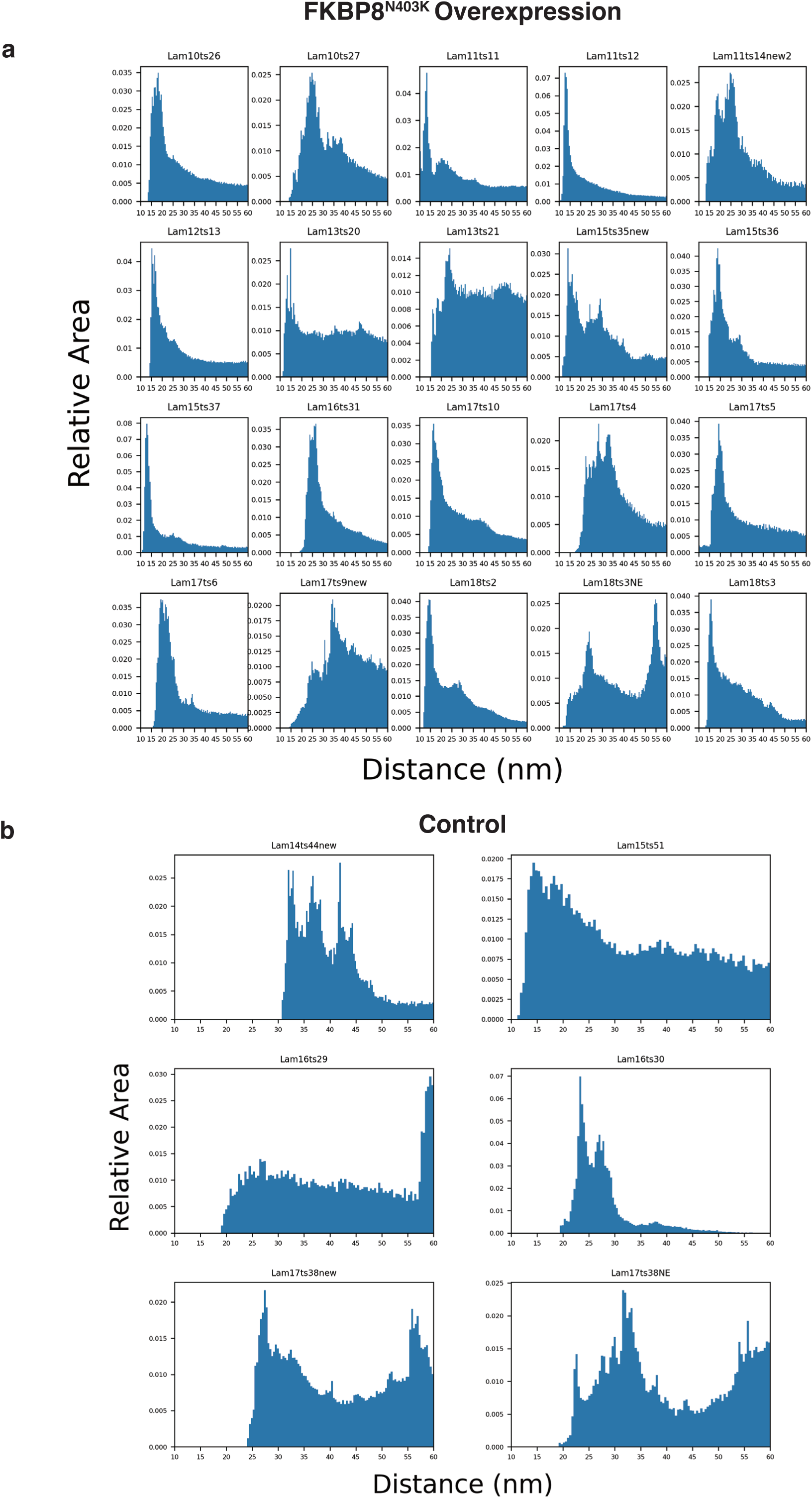
All ER-OMM distance histograms used for the aggregate analysis in Figure 7. **a, b,** All ER-OMM distance histograms for FKBP8^N403K^ OE cells (**a**) and the control cells (**b**) used in Fig. 7g

**Extended Data Fig 10.**
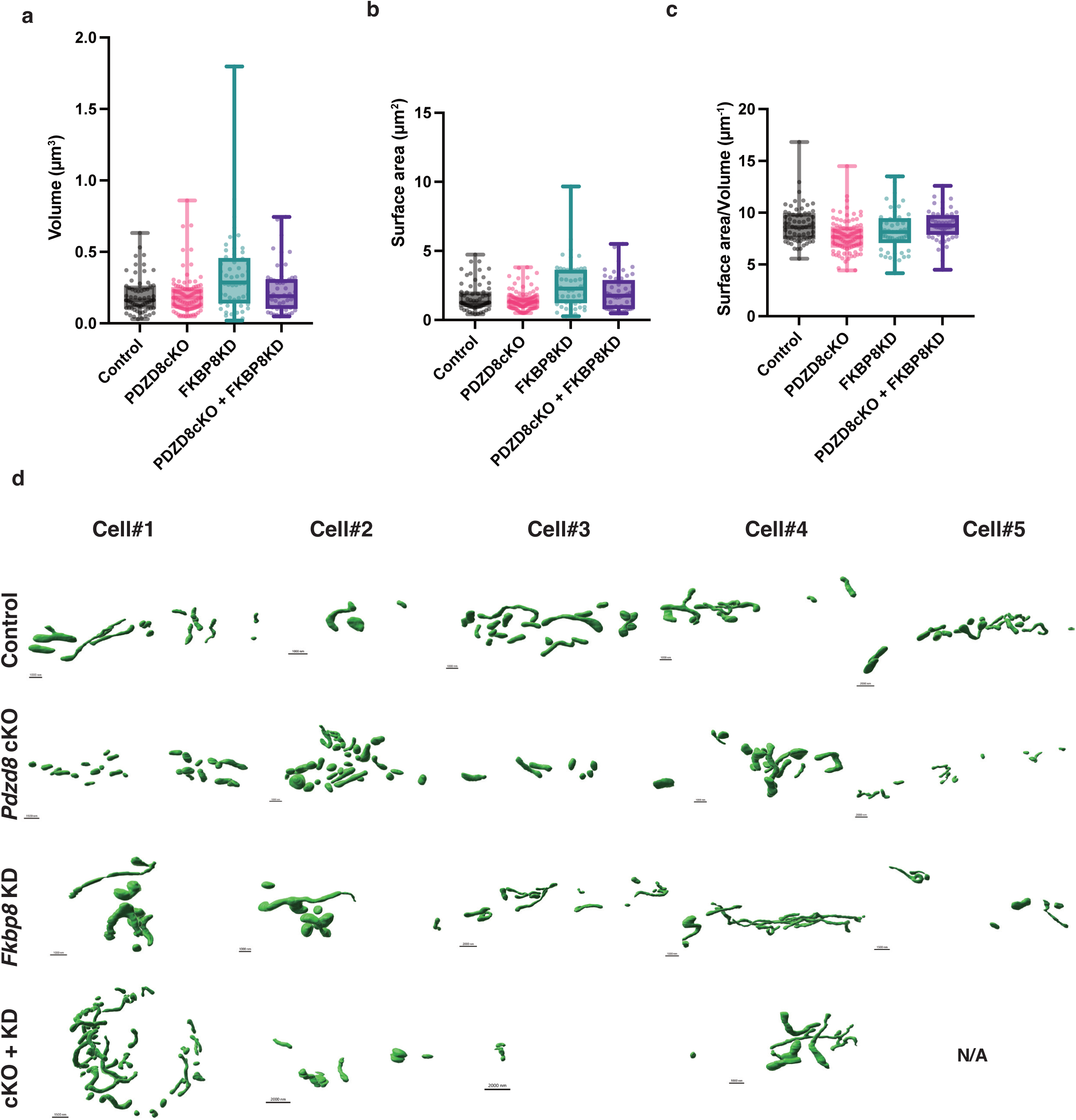
Morphological properties of mitochondria analyzed by array-tomography. **a-c,** Quantifications of the volume (**a**), surface area (**b**), and surface/volume ratio (**c**) of individual mitochondria in the control, *Pdzd8* cKO, *Fkbp8rd* KD, and *Pdzd8* cKO + *Fkbp8* KD cells analyzed in Fig. 8c. **c,** All the 3D reconstructions of mitochondria analyzed in this study.

